# Epitranscriptome changes triggered by ammonium nutrition regulate the proteome response of maritime pine roots

**DOI:** 10.1101/2021.04.20.440618

**Authors:** Francisco Ortigosa, César Lobato-Fernández, Juan Antonio Pérez-Claros, Francisco R. Cantón, Concepción Ávila, Francisco M. Cánovas, Rafael A. Cañas

## Abstract

Epitranscriptomic modifications constitute a gene expression checkpoint in all living organisms. As nitrogen is an essential element for plant growth and development, a reasonable hypothesis is that changes in the epitranscriptome may regulate nitrogen acquisition and metabolism. In this study, epitranscriptomic modifications caused by ammonium nutrition were monitored in maritime pine roots through direct RNA sequencing using Oxford Nanopore Technology. Transcriptomic responses mainly affected transcripts involved in nitrogen and carbon metabolism, defense, hormone synthesis/signaling, and translation. Global detection of epitranscriptomic marks was performed to evaluate this posttranscriptional mechanism in untreated and ammonium-treated seedlings. Increased m^6^A deposition in the 3’-UTR was observed in response to ammonium, which seems to be correlated with poly(A) lengths and changes in the relative abundance of the corresponding proteins. The results showed that m^6^A deposition and its dynamics seem to be important regulators of translation under ammonium nutrition. These findings suggest that protein translation is finely regulated through epitranscriptomic marks likely by changes in mRNA poly(A) length, transcript abundance and ribosome protein composition. An integration of multiomics data suggests that the epitranscriptome modulates responses to developmental and environmental changes, including ammonium nutrition, through buffering, filtering, and focusing the final products of gene expression.

## INTRODUCTION

Discoveries in the nascent field of molecular biology culminated in the central dogma of molecular biology during the 1970s (Crick, 1970). Currently, it is known that transcription and translation processes are not always directly linked. Multiple factors intervene in the gene response and its regulation. Two good examples of this are long noncoding RNAs (lncRNAs) (Liu et al., 2015) and microRNAs (miRNAs) (Paul et al., 2015). These types of RNA regulate important aspects of both development and response to external stimuli in plants (Liu et al., 2015; Paul et al., 2015). However, these studies represent only some aspects of overall RNA metabolism. In recent years, an increasing interest has focused on determining the biological role of RNA chemical modifications emerging as a new field of study under the term *epitranscriptomics*. These modifications are found in all RNA types, such as transfer RNAs (tRNAs), ribosomal RNAs (rRNAs), messenger RNAs (mRNAs) and small RNAs (RNAs) (Xiong et al., 2017). To date, more than 160 different modifications have been identified in RNA (Shen et al., 2019). In *Arabidopsis thaliana*, m^7^G, m^6^A, m^1^A, m^5^C, hm^5^C, and uridylation have been identified as modifications in mRNA (Shen et al., 2019). The marriage between classical detection techniques and high-throughput sequencing has allowed to determine N^6^-methyladenosine (m^6^A) as the most prevalent chemical modification in mRNAs, both in animals and plants (Fray and Simpson, 2015). Transcriptome-wide analysis revealed that the m^6^A mark in transcripts is predominantly localized near the stop codon and throughout the 3′ untranslated region (UTR) (Shen et al., 2019). An m^6^A methylation motif (RR**A**CH [R = A/G; H = A/U/C; **A** = m^6^A]) that is conserved between plants and other eukaryotic organisms has been described (Shen *et al.,* 2019). m^6^A deposition, recognition and elimination are carried out by different proteins (Shen et al., 2019). Several cellular functions have been observed to be affected by m^6^A modification, such as mRNA stability (Wei et al., 2018) or translational efficiency (Luo et al., 2014). In addition, proper m^6^A deposition has been reported to be essential during *Arabidopsis* embryo development (Zhong et al., 2008) and to take part in biotic and abiotic plant stress responses (Martínez-Pérez et al., 2017; Anderson et al., 2018), fruit ripening (Zhou et al., 2019), flowering transition (Duan et al., 2017), leaf morphogenesis (Arribas-Hernández et al., 2018), trichome development (Bodi et al., 2012) and apical shoot meristem development (Shen et al., 2016).

Nevertheless, little is known about the potential role of the epitranscriptome in the regulation of nitrogen (N) nutrition, with only the involvement of m^6^A in the regulation of nitrate assimilation and signaling having been studied (Hou et al., 2021). However, N is an essential element for plant growth and development and a key component of cellular constituents such as nucleic acids, proteins, and chlorophylls (Hawkesford et al., 2012). In soils, plants can take up N inorganic forms such as nitrate (NO_3_^-^) or ammonium (NH_4_^+^) and organic forms such as urea, peptides, or amino acids (Jia and von Wirén, 2020). Plants such as rice, tea or maritime pine prefer NH_4_^+^ over NO_3_^-^ as the main N source (Sasakawa and Yamamoto, 1978; Ruan et al., 2016; Ortigosa et al., 2020). In many plants, important changes in the transcriptome, proteome and metabolome have been described in relation to nitrogen nutrition and have mainly focused on the supply of NO_3_^-^ and NH_4_^+^ (Patterson et al., 2010; Yang et al., 2018; Ravazzolo et al., 2020). In this way, it has been observed that these N forms trigger both shared and differential responses involving different pathways and many result in phenotypic differences such as specific changes in the root system architecture (RSA) and growth (Jia and von Wirén, 2020). Therefore, it is reasonable to hypothesize that some of these described responses to N nutrition may be influenced by epitranscriptome regulatory processes.

Oxford Nanopore Technology (ONT) is a third-generation sequencing platform that is currently the only option for direct sequencing of RNA samples without the requirement of reverse transcription and amplification steps (Parker et al., 2020). These characteristics are of great relevance in transcriptomics since they allow the reduction of sequencing biases and maintenance of nucleoside modifications that facilitate epitranscriptomics studies (Gao et al., 2021).

The aim of the present work is to shed light on the short-term response of maritime pine roots to NH_4_^+^ nutrition, elucidating what kind of regulatory relationship exists between transcriptomics, epitranscriptomics, proteomics and metabolite profiles. For this purpose, cutting edge and commonly used omics approaches, such as comprehensive transcriptome analyses by direct RNA sequencing (DRS) using Oxford Nanopore Technology (ONT), epitranscriptomic modification detection focused on m^6^A assisted by the ONT platform, and quantitative proteomic and metabolite profiling, were combined in the present study.

## RESULTS

### Metabolite profiling in response to NH_4_^+^ nutrition

Changes in the amounts of polar metabolites caused by short-term NH_4_ nutrition were analyzed in maritime pine roots (Supplemental Dataset 1). A statistically significant decrease in the amounts of carbohydrates such as sucrose and glucose was observed (Figure 1; Supplemental Figure 1). Additionally, a statistically significant increase in shikimate levels was noted. In turn, although the results were not statistically significant, there was a slight trend of increased levels of glycolysis intermediary metabolites such as glucose-6 phosphate and glucose-1 phosphate as well as TCA anaplerotic pathway intermediates such as cis-aconitate and GABA. In parallel, an increase in the accumulation of L-glutamine, L-glutamate, and L-methionine, as well as in aromatic amino acids (L-phenylalanine and L-tryptophan), was observed in NH_4_^+^-treated plants. In contrast, metabolites such as L-asparagine, isocitrate, 2-oxoglutarate and L-malate were observed to decrease.

**Figure 1.**
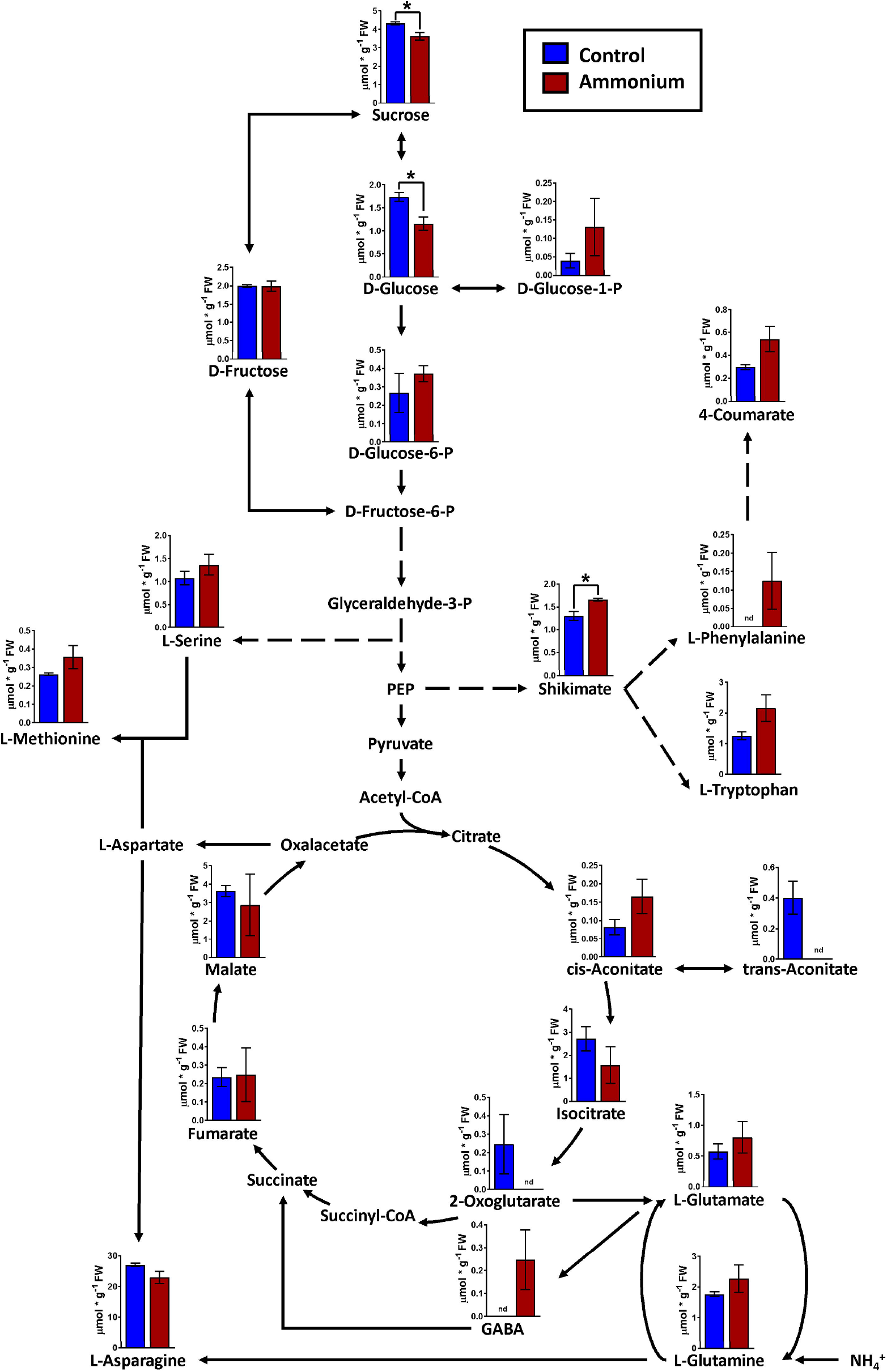
Main trend of metabolites in the roots of control and NH_4_^+^-treated seedlings at 24 hours post-irrigation. Blue columns correspond to control plants; red columns correspond to 3 mM NH_4_^+^ supply. Differences between treatments were determined with a t-test. Significant differences are indicated with asterisks on top of the columns: * at *P* < 0.05. Error bars show SE with *n* = 3. PEP: phosphoenolpyruvate.

### Direct RNA sequencing (DRS)

The global sequencing results are shown in Supplemental Table 1. The mean read sizes were between 908 bp to 1059 bp (Figure 2A). The longest reads ranged from 10298 bp to 14299 bp. Differential expression analyses identified 350 differentially expressed (DE) transcripts. From the total DE transcripts obtained, 106 were upregulated and 244 were downregulated (Figure 2B; Supplemental Dataset 2). Singular enrichment analysis (SEA) was performed individually for each gene expression regulation (up- and downregulated) to classify the biological functions under NH_4_^+^ nutrition. The SEA global results are shown in Supplemental Dataset 3. Upregulated transcripts were significantly enriched with GO terms for Biological Process (BP) such as “ammonia assimilation cycle” (GO:0019676), “protein glutathionylation” (GO:0010731), “response to cadmium ion” (GO:0046686) and “developmental growth” (GO:0048589); for Cellular Component (CC) such as “chloroplast stroma” (GO:0009570) and “cytosolic ribosome” (GO:0022626); and for Molecular Function (MF) such as “glutamate synthase (NADH) activity” (GO:0016040), “acylglycerol lipase activity” (GO:0047372) and “phospholipase activity” (GO:0004620) (Figure 2C; Supplemental Figure 2). The downregulated transcripts were enriched in the BP terms “response to heat” (GO:0009408), “response to water deprivation” (GO:0009414), “protein folding” (GO:0006457) and “ethylene-activated signaling pathway” (GO:0009873) and the MF terms “calcium ion binding” (GO:0005509), “transcription coactivator activity” (GO:0003713), “mRNA binding” (GO:0003729), “chaperone binding” (GO:0051087) and “transcription factor activity, sequence-specific DNA binding” (GO:0003700). A more detailed study of the upregulated transcriptomic response revealed that *PpGS1b* (pp_68481) and *PpNADH-GOGAT* (pp_238920) were upregulated. Interestingly, transcripts for genes involved in defense-related response were upregulated: *PpAMP1* (pp_58005, pp_58008), different class IV chitinases (pp_239593, pp_239598, pp_239600, pp_117809), different splicing coding forms of patatin-like protein 2 (pp_71017, pp_71018, pp_71019, pp_71020, pp_71022), a PR-1 pathogenesis-related protein (pp_87427), a defensin coding transcript (pp_92119) and an RPW8 domain-containing protein (pp_142311) (Supplemental Dataset 2). Cell wall-related transcripts were also upregulated, such as those encoding expansin-A18 (pp_134987), nonclassical arabinogalactan protein 30 (pp_66323), probable prolyl 4-hydroxylase 4 (pp_235715), xyloglucan:xyloglucosyl transferase (pp_68519) and different forms of xyloglucan endotransglucosylase/hydrolase (pp_66707, pp_66708, pp_68517), was also observed. In contrast, the downregulation of different transcription factors (TFs) was observed (Supplemental Dataset 2), such as ethylene response factors (ERFs) (pp_58625, pp_58626, pp_86737, pp_96228, pp_96234), a trihelix transcription factor (pp_59947), and a MYB coding transcript (pp_202778). The repression of different splicing forms of an auxin-repressed protein/dormancy-auxin associated protein coding transcript (pp_58457, pp_58458, pp_58459, pp_58461, pp_58462, pp_58463), the SnRK1 regulator FCS-like zinc finger transcript (pp_202096), and transcripts encoding carbohydrate metabolism enzymes such as pyruvate decarboxylase (pp_78343, pp_123611, pp_126347) and sucrose synthase (pp_144843) was also observed. The differential expressions observed in DRS was confirmed by RT-qPCR analyses of seven transcripts including *SAM synthase, MYB5, PFK, AMP1, NADH-GOGAT, GS1b,* and *ASPG*. The results showed the same trend between the DRS and RT-qPCR results for all DE transcripts, with some differences in the logFC values (Figure 2D).

**Figure 2.**
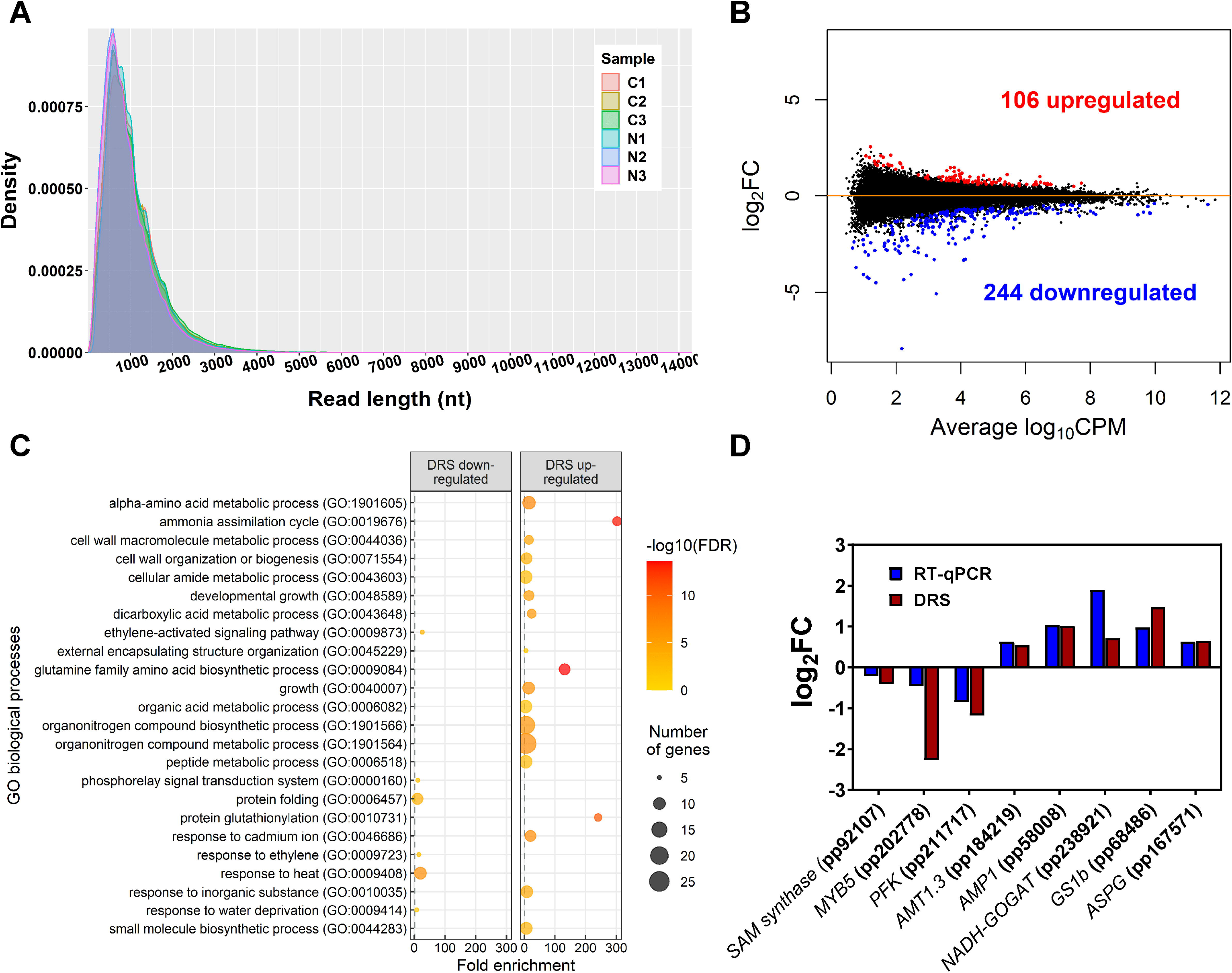
Main DRS transcriptomics results. **A.** Size distribution of reads obtained through DRS-ONT sequencing. **B.** MA plot containing all detected transcripts. Blue points correspond to significantly downregulated transcripts in maritime pine roots at 24 h after the treatment with 3 mM ammonium. Red points correspond to significantly upregulated transcripts in maritime pine roots at 24 h after the treatment with 3 mM ammonium. **C.** Significant GO terms from significant DE transcripts after a SEA analysis. **D.** Comparison between DRS and RT-qPCR results from different transcripts with significant differential expression in the DRS analysis.

### Differential DRS epitranscriptomics

To determine the differential epitranscriptomic marks, DRS results were explored to identify the chemically modified nucleosides in the mRNAs using Tombo software. After analysis, 513 statistically significant putative modified nucleosides were obtained in 283 transcripts (Figure 3A; Supplemental Dataset 4). There were 221 undermodified positions in 161 transcripts and 292 overmodified positions in 184 transcripts. Among them, 58 transcripts had significant over- and undermodified positions, including *PpGS1b* (pp_68474) and *translationally-controlled tumor protein* (*TCTP*, pp_72505) (Supplemental Dataset 5). The percentage of modifications for each kind of nucleoside was similar for adenosine, guanosine, and uridine (28%, 29% and 25%) but lower for cytidine (17%), without any obvious difference between sample conditions (Figure 3B). When the global set of modification ratios and transcript amounts were compared (Figure 3C), significant and negative correlations were found to be slightly stronger under NH_4_^+^ supply (−0.36) than under the control (−0.30) (Figure 3C). Modification ratios of individual nucleosides compared to transcript amounts were also similar (Supplemental Figure 3).

**Figure 3.**
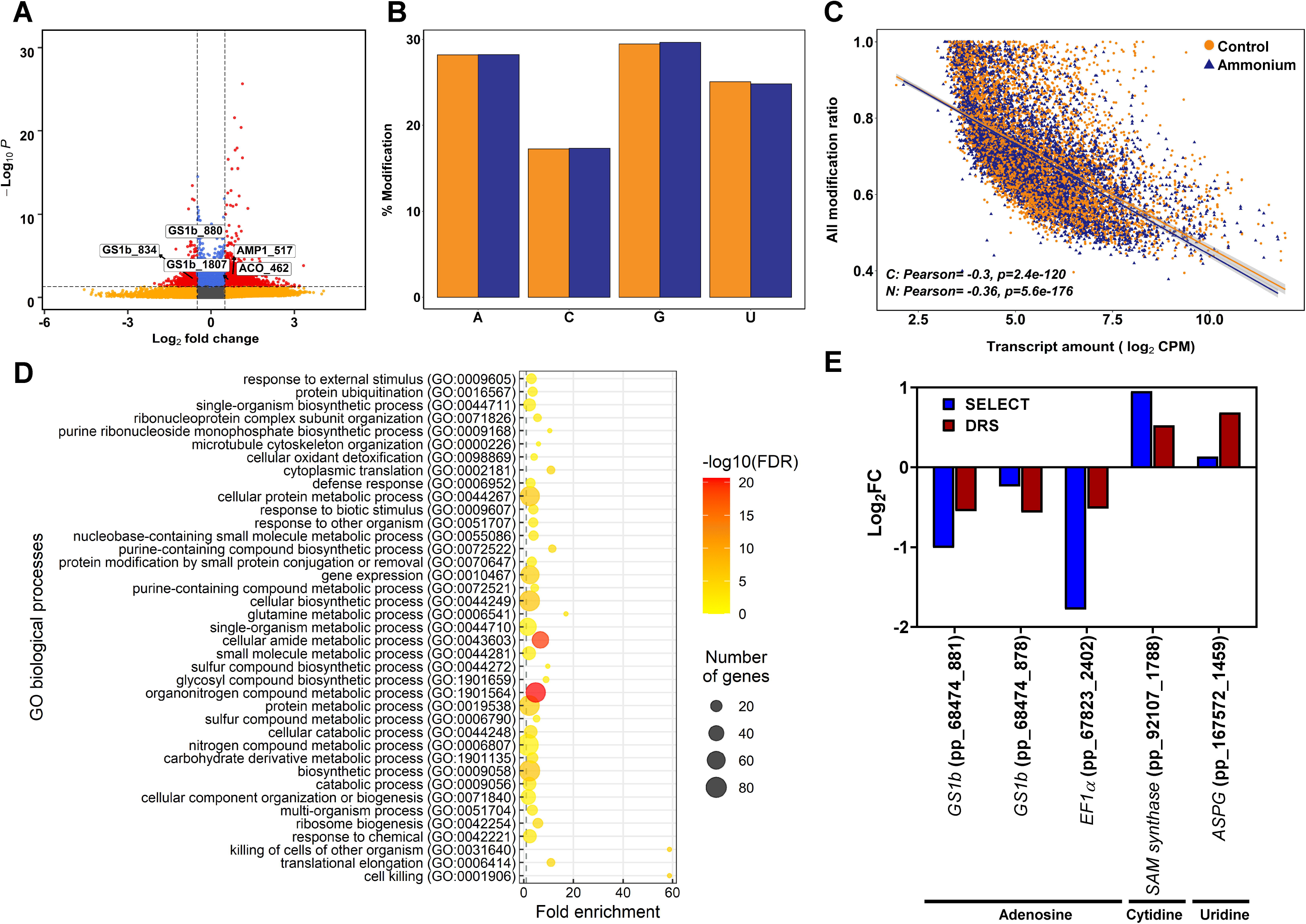
Epitranscriptomics modification results from Tombo software analysis. **A.** Volcano plot of the detected epitranscriptomics modifications. Yellow points correspond to modifications with *P*-values lower than 0.05 but with an absolute logFC value lower than 0.5. Blue points correspond to modifications with an absolute logFC value higher than 0.5 but with *P*-values higher than 0.05. Red points correspond to significant DE modifications with an absolute logFC value higher than 0.5 and *P*-values lower than 0.05. **B.** Percentage of detected modifications per nucleobase. Yellow columns correspond to control samples. Purple columns correspond to 3 mM ammonium treated samples. **C.** Scatter plot and correlation between detected modifications using Tombo software and transcript amounts determined by DRS sequencing. **D.** Significant GO terms from significant DE modifications after a SEA analysis. **E.** Comparison between DRS and SELECT results from different transcript modified positions with significant differential modification in the Tombo analysis.

The functions of the transcripts with DE modifications were analyzed using SEA (Figure 3D; Supplemental Figure 4; Supplemental Dataset 6). A total of 116 significant GO terms were obtained. At the BP level, the main functions were related to the terms “ribosome biogenesis” (GO: 0042254); “translation” (GO:0006412), “proteolysis” (GO:0006508), “protein ubiquitination” (GO:0016567), “response to stress” (GO:0006950), “response to external biotic stimulus” (GO:0043207) and “glutamine metabolic process” (GO:0006541). Similarly, the “ribosome” term (GO:0005840) was the main function at the CC level. Finally, the terms “GTPase activity” (GO:0003924), “mRNA binding” (GO:0003729), “translation factor activity, RNA binding” (GO:0008135), “ubiquitin protein ligase binding” (GO:0031625), “ubiquitin conjugating enzyme activity” (GO:0061631) and “endopeptidase inhibitor activity” (GO:0004866) were the main functions at the MF level.

The differential modifications were verified using SELECT. This qPCR-based technique was initially designed to determine differential deposition of m^6^A in total RNA mixtures. However, nucleoside modifications putatively detected in cytidine and uridine by Tombo were also detected with SELECT (Figure 3E). The obtained results for each position showed a similar trend between the differential epitranscriptomic results from SELECT and Tombo (Figure 3E).

### m^6^A identification

The bioinformatic pipeline Nanom6A was used to specifically identify m^6^A modifications in the RRACH sites from DRS data (Supplemental Dataset 7). The distribution of m^6^A sites along the full-length transcripts showed a higher accumulation of marks over two-thirds of the relative length in control samples while a higher accumulation of marks in the final portion of the transcripts was observed in NH_4_^+^-treated samples (Figure 4A). A more detailed distribution study showed that m^6^A sites were more abundant in the 5’-UTR and coding (CDS) regions under the control condition, while m^6^A sites tended to be more abundant in the 3’-UTR regions of transcripts isolated from NH_4_^+^-treated seedlings (Figure 4B). The m^6^A frequency was higher in the CDS, mainly in the central portion, and at the beginning of the 3’-UTR regions, but it was lower at the transcript ends (Figures 4A and 4B).

**Figure 4.**
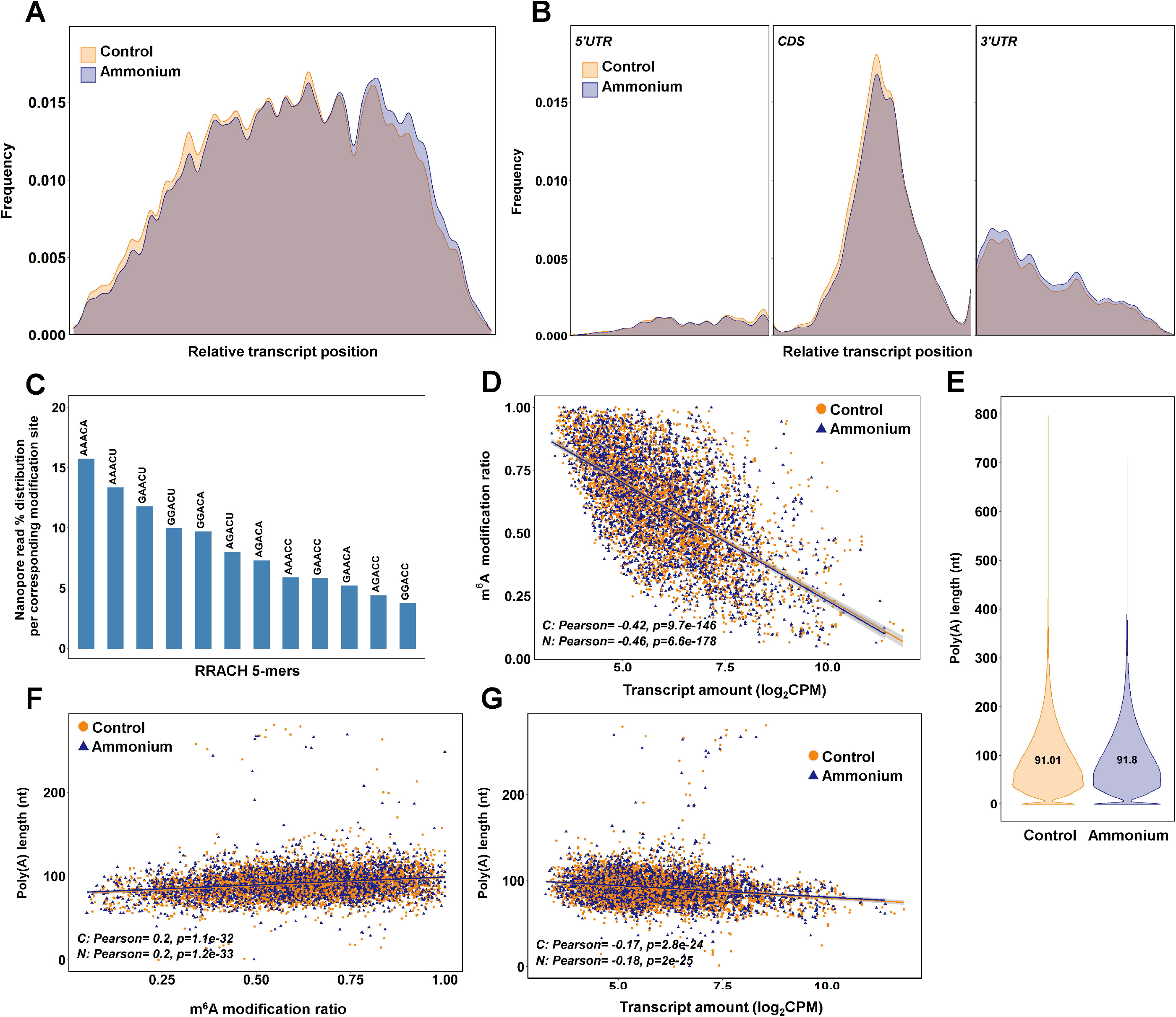
m^6^A identification results using the bioinformatics pipeline Nanom6A. **A.** Distribution of the identified m^6^A all along the transcripts. **B.** Distribution of the identified m^6^A along each part of the transcripts: 5’UTR, CDS and 3’UTR respectively. **C.** Percentage of identified m^6^A for each different variant (kmer) of the consensus sequence RRACH for adenine methylation. **D.** Scatter plot and correlations between m^6^A modification ratio and transcript amount from DRS sequencing. **E.** Poly(A) length of the transcripts identified in the DRS sequencing. **F.** Scatter plot and correlations between poly(A) length and m^6^A modification ratio. **G.** Scatter plot and correlations between poly(A) length and transcript amount from DRS sequencing.

The most predominant RRACH sequence was AAACA (>15%), while GGACC was the least abundant (< 5%) (Figure 4C). The comparison between m^6^A modification ratios and transcript amounts showed significant negative correlations for the control (−0.42) and NH_4_ conditions (−0.46) (Figure 4D). The lengths of the poly(A) tails were determined from DRS data (Figure 4E). As expected, most of the poly(A) tails had a size between 40-250 nt with similar means in both conditions, 91.01 nt and 91.8 nt. The poly(A) tail lengths had significant positive correlations with m^6^A ratios in both conditions (0.2) (Figure 4F). Significant but lower positive correlations were found when poly(A) tail lengths were compared with global and nucleoside modification ratios obtained with Tombo (Supplemental Figure 5). Finally, the poly(A) tail lengths and transcript amounts exhibited significant negative correlations for the control (−0.17) and NH_4_^+^ conditions (−0.18) (Figure 4G).

### Differential proteomics

A total of 2,385 proteins were identified in the shotgun proteomics analysis (Supplemental Dataset 8). Among the identified proteins, 114 were differentially regulated by NH_4_^+^: 38 were more abundant, while 76 were less represented (Figure 5A; Supplemental Dataset 8). To elucidate the biological roles of the identified proteins, SEA analyses were performed (Figure 5B; Supplemental Figure 6; Supplemental Dataset 9). The upregulated proteins showed as representative BP terms “cell redox homeostasis” (GO:0045454), “protein complex assembly” (GO:0006461), “translation” (GO:0006412), “cellular response to oxidative stress” (GO:0006979) and “defense response to bacterium” (GO:0042742). At CC level, “ribosome” (GO:0005840), “nucleolus” (GO:0005730) and “chloroplast” (GO:0009507) were the enriched terms. Finally, “structural constituent of ribosome” (GO:0003735) and “enzyme regulator activity” (GO:0030234) were the significant MF terms. Among the downregulated proteins, the representative enriched functions were “ribosome assembly” (GO:0042255), “ATP metabolic process” (GO:0046034), “translation” (GO:0006412), “response to metal ion” (GO:0010038) and “oxidation-reduction process” (GO:0055114) among the BP terms. At the CC level, the significant terms were “cytosol” (GO:0005829), “apoplast” (GO:0048046), “chloroplast” (GO:0009507) and “ribosome” (GO:0005840). “Oxidoreductase activity, acting on the aldehyde or oxo group of donors, NAD or NADP as acceptor” (GO:0016620), “structural constituent of ribosome” (GO:0003735), “GTPase activity” (GO:0003924) and “GTP binding” (GO:0005525) were the enriched MF terms.

**Figure 5.**
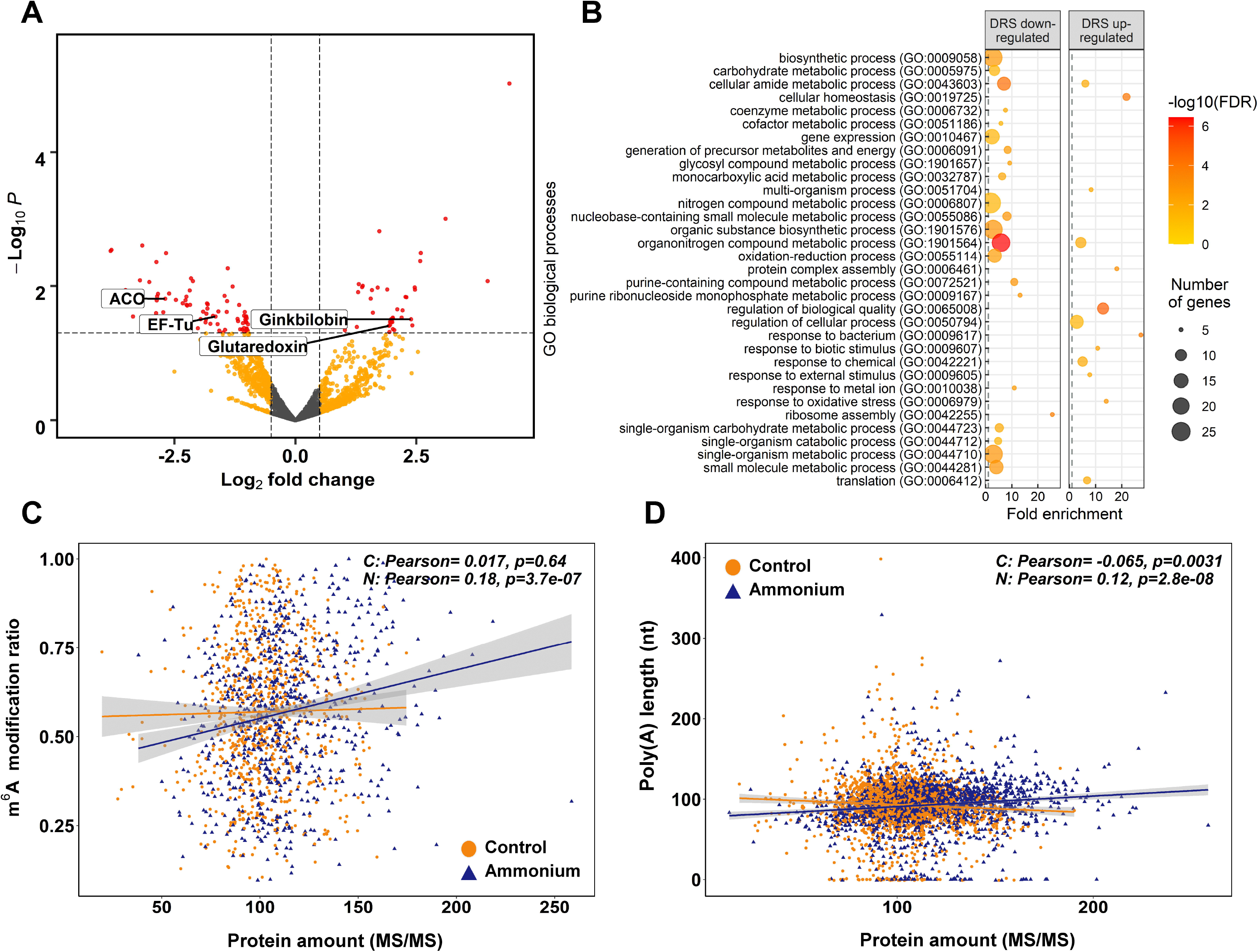
Differential expression proteomics results. **A.** Volcano plot of the identified proteins. Yellow points correspond to modifications with *P*-values lower than 0.05 but with an absolute logFC value lower than 0.5. Red points correspond to significant DE modifications with an absolute logFC value higher than 0.5 and *P*-values lower than 0.05. **B.** Significant GO terms from significant DE proteins after a SEA analysis. **C.** Scatter plot and correlations between m^6^A modification ratios and protein amounts. **D.** Scatter plot and correlations between transcript amounts from DRS sequencing and protein amounts.

The putative relationship between the m^6^A modification ratio and protein abundance was determined through Pearson correlation analysis (Figure 5C). In NH_4_^+^-treated seedlings, a significant positive correlation was observed (0.18), while there was no correlation in control seedlings. The same effect was observed between nucleoside modification ratios from Tombo and protein amounts (Supplemental Figure 7). In addition, similar correlation results were obtained when poly(A) tail lengths and protein amounts were compared, and only NH_4_^+^-treated roots had a significant positive correlation (0.12) (Figure 5D).

### Integration of data from omics approaches

The results of the different omics approaches employed in the present work were integrated to explore possible regulatory steps in response to NH_4_^+^ supply in maritime pine. The global results showed 30 different elements/genes with significant results based on at least two approaches at least; 14 of them were significant in DRS and epitranscriptomics, 5 in DRS and proteomics, 9 in epitranscriptomics and proteomics, and 2 in all approaches (Figure 6A; Supplemental Dataset 10). Altogether, the genes/proteins identified were involved in N metabolism (*PpASPG*, *PpGS1b,* alanine-glyoxylate aminotransferase and isocitrate dehydrogenase), defense (*PpAMP1*, a ginkbilobin and a chitinase), oxidative stress response (alcohol dehydrogenase, aldehyde dehydrogenase, peroxidase and glutaredoxin), translation (ribosomal proteins and elongation factors) and RNA binding (cold shock proteins and a BURP domain protein RD22). Among them, the presence of the 1-aminocyclopropane-1-carboxylate (ACC) oxidase, which had a putative modification at the position 462 on the contig, must be highlighted. This epitranscriptomic mark was overexpressed (logFC 0.84) under NH_4_^+^ nutrition, while the ACC oxidase protein was underexpressed (logFC −2.69). Similarly, changes in transcript level, protein level and transcript modification ratio can show opposite trends with variations between genes. For the aldehyde dehydrogenase, the transcript expression and modification were overexpressed (0.58 and 0.73, respectively), but protein expression was underexpressed (logFC −1.20). However, for the CSP/GRP (pp_211512 and pp_211516), transcript accumulation was underexpressed (−0.43 and −0.7) and the epitranscriptomic modifications and protein expression were overexpressed (0.88-0.73 and 2.11 respectively).

**Figure 6.**
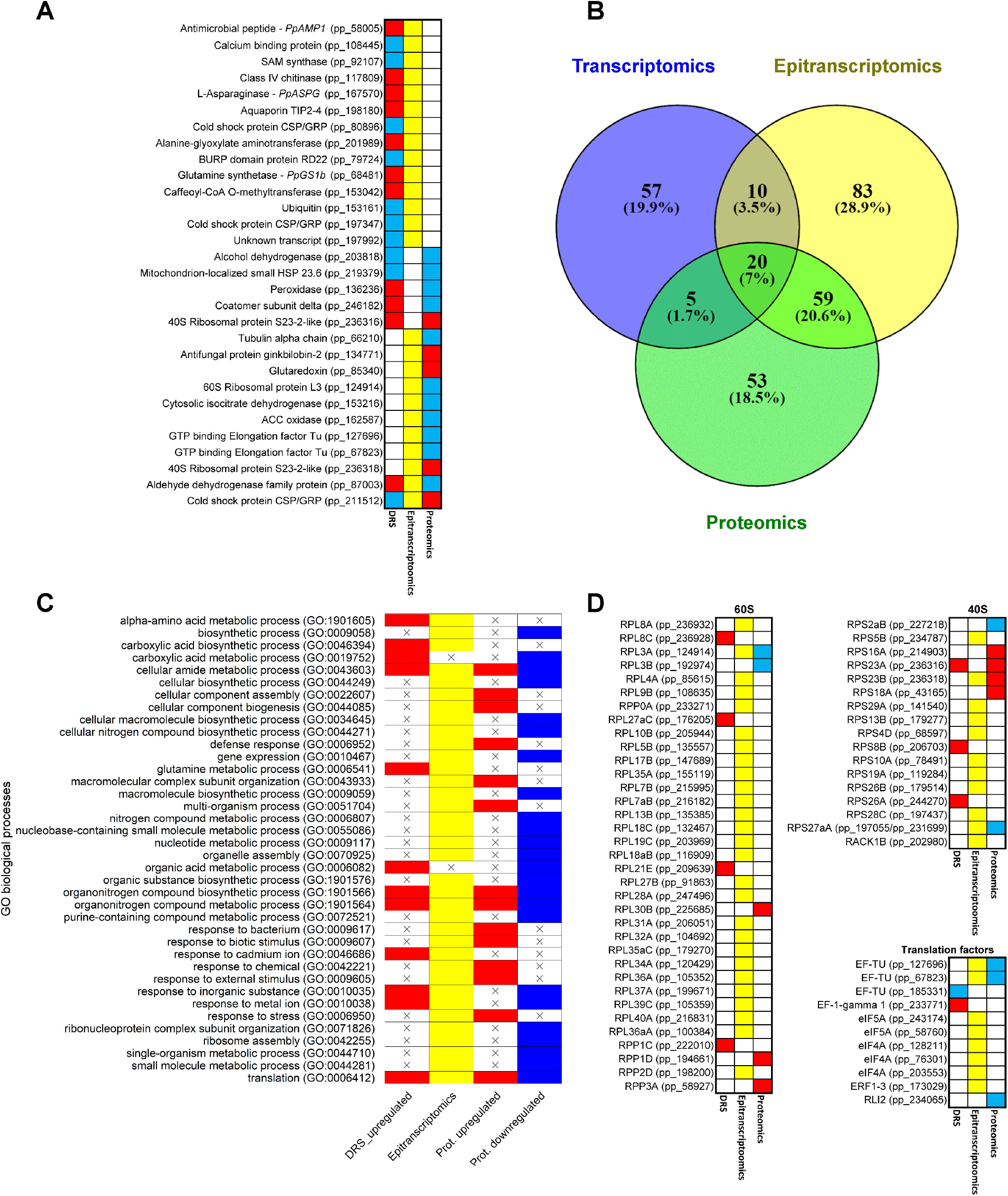
Common results among different omics approaches. **A.** Genes with significant expressions in more than one omics approach. Red boxes correspond to upregulated transcripts and proteins. Blue boxes correspond to downregulated transcripts and proteins. Yellow boxes correspond to significant differentially modified positions in the transcripts. **B.** Venn diagram of significant GO terms in the different omics approaches. **C.** Heatmap of significant GO terms shared by at least two omics approaches. **D.** Heatmap of genes that produce proteins involved in ribosome and translation. Ribosomal protein equivalences obtained from Martinez-Seidel et al. (2020). Red boxes correspond to upregulated transcripts and proteins. Blue boxes correspond to downregulated transcripts and proteins. Yellow boxes correspond to significant differentially modified positions in the transcripts.

The comparison between the significant GO terms in the omics approaches revealed that 94 of the 287 GO terms were shared (Figure 6B-6C; Supplemental Dataset 11). Interestingly, 20 (7%) of them were common to the three sets of results. The epitranscriptomics and proteomics comparison included the highest number of GO terms (59, 20.6%), and the transcriptomics and proteomics comparison included the lowest number, with only 5 (1.7%). The most representative GO terms common to the three omics datasets were “ribosome” (GO:0005840), “structural constituent of ribosome” (GO:0003735) and “translation” (GO:0006412). Between the transcriptomics and epitranscriptomics datasets, “mRNA binding” (GO:0003729) and “glutamine metabolic process” (GO:0006541) were the main GO terms. In the transcriptomics and proteomics comparison, “oxoacid metabolic process” (GO:0043436) was the main GO term. Finally, between epitranscriptomics and proteomics the main GO terms were “small ribosomal subunit” (GO:0015935), “GTPase activity” (GO:0003924), “ribosome assembly” (GO:0042255), “defense response” (GO:0006952) and “response to external stimulus” (GO:0009605). According to these functional results, several transcripts coding for eukaryotic ribosomal proteins (38) and translation factors (8) had differential epitranscriptomic marks based on the Tombo results (Figure 6D).

## DISCUSSION

The response of maritime pine roots to NH_4_^+^ nutrition has been studied from a multiomics perspective that includes direct RNA sequencing using the ONT platform, which has allowed a global epitranscriptomics analysis. Although N is an essential nutrient for proper plant growth and development (Xu et al., 2012), little is known about the role of epitranscriptomic marks in the regulation of gene expression in response to N nutrition. The results in the present work highlight the importance of epitranscriptomic marks in the regulation of gene expression.

### Epitranscriptome changes in response to NH_4_^+^nutrition

Correlation of global epitranscriptomics results with transcript abundance (Figure 3C), especially m^6^A modifications (Figure 4D), are consistent with those from previous works in *Populus* (Gao et al., 2021) and *Arabidopsis* (Luo et al., 2014; Wan et al., 2015) and support the role of m^6^A in mRNA turnover, as previously described in mammals (Wang et al., 2014; Li et al., 2019). Additionally, m^6^A identification revealed that in the roots of maritime pine, AAACA and AAACU were the most abundant RRACH 5-mer motifs (Figure 4C), as previously reported in *Arabidopsis*, maize, and poplar (Wan et al., 2015; Miao et al., 2020; Parker et al., 2020; Gao et al., 2021). These findings strongly suggest conservation of the RNA m^6^A methylation machinery in plants. However, pine m^6^A distribution, with similar levels of enrichment in the middle of the CDS and at beginning of the 3’-UTR (Figure 4A-4B), slightly differs from the obtained results in angiosperms, where a greater enrichment of m^6^A was observed in the 3’-UTR (Miao et al., 2020; Parker et al., 2020; Gao et al., 2021). This discrepancy may be attributed to the lack of a well-curated reference genome for maritime pine. Interestingly, the distribution of m^6^A sites in pine transcripts is dynamically regulated by increasing the m^6^A deposition in the 3’-UTR in response to NH_4_^+^, which seems to be correlated with poly(A) length and changes in protein abundance (Figure 6C-6D), as well as the identification of transcripts with upregulated and downregulated epitranscriptomic marks (Supplemental Dataset 5). This observation agrees with previous results in other eukaryotic organisms (Meyer et al., 2012; Anderson et al., 2018).

A previous work in *Caenorhabditis elegans* reported that highly expressed transcripts contained a relatively short and well-defined poly(A) tail (Lima et al., 2017), which has been related to translational efficiency (Wu et al., 2020). Our results reveal that increased poly(A) tail lengths correlated with higher m^6^A and lower transcript abundances (Figure 4F-4G). These findings are consistent with the effect of m^6^A modifications on transcript levels (Figure 4D), which follow the same trend as previously published results (Luo et al., 2014; Wan et al., 2015; Parker et al., 2020; Gao et al., 2021). The integration of all these results with the positive correlations of m^6^A ratios and poly(A) tail length with protein amounts (Figure 5C-5D) suggests that m^6^A could affect in the translation efficiency of the differentially modified transcripts, as previously described in mammals (Meyer, 2019). This evidence could also explain, at least in part, the lack of a relationship between the transcriptomic, proteomic and metabolite data.

Epitranscriptomic marks, mainly m^6^A, seem to modulate protein synthesis through mRNA stability and modify the translation processes. This is in line with a previous hypothesis considering that initial gene responses caused by environmental factors are modulated at the level of final products of gene expression to maintain cellular homeostasis (Cañas et al., 2015). Thus, the variable transcriptomic response can be controlled through intermediate steps, such as epitranscriptomic regulation, to generate an appropriate and stable cellular response.

### Carbon and nitrogen metabolism

As expected, N assimilation was induced by NH_4_^+^, as shown by the significant increase in *GS1b* and *NADH-GOGAT* transcripts and the concomitant accumulation of L-glutamine and L-glutamate (Figure 1 and 7; Supplemental Dataset 3), consistent with previous transcriptomic reports (Li et al., 2017; Sun et al., 2017). However, no upregulation of GS1b and NADH-GOGAT was observed at the proteomic level (Supplemental Dataset 8), even though a trend towards higher GS activity was observed in the presence of NH_4_^+^ (Supplemental Figure 8). Similar proteomics results for NH_4_^+^ nutrition have been described before (Marino et al., 2016; Coleto et al., 2019), as well as the lack of correlation between relative expression levels of GS transcripts and GS activity. To explain these discrepancies, it was hypothesized that the response to N nutrition could be regulated through a posttranscriptional (translational or posttranslational) mechanism (Ortigosa et al., 2020). In this way, several nucleosides with differential epitranscriptomic marks, including m^6^A, have been identified in *GS1b* transcripts, which could explain the differences between transcriptomics and proteomics data (Supplemental Dataset 4 and 7). Accordingly, in *Arabidopsis*, m^6^A can regulate the alternative polyadenylation and transcript abundance of the *GLN1;1* and *GLN1;3* genes during nitrate assimilation and signaling (Hou et al., 2021).

Additionally, the results of this study suggest the existence of a carbon flux through glycolysis and the TCA cycle to provide carbon skeletons for N assimilation (de la Peña et al., 2019). Interestingly, the fermentation pathway seems to be repressed, as suggested by the downregulation of pyruvate decarboxylase (PDC) transcripts and the decrease in alcohol dehydrogenase (ADH) and aldehyde dehydrogenase (ALDH) protein levels (Supplemental Datasets 2 and 7). It is well known that plants under low oxygen conditions regulate their metabolism, inducing the fermentation pathway in which PDC and ADH activities are essential, which results in pyruvate consumption, the production of ethanol and the concomitant oxidation of NADH to NAD^+^ (Mithran et al., 2014). The repression of the fermentative pathway and the overrepresentation of TIM proteins (inner membrane translocase proteins) (Supplemental Dataset 7) highlight the need for organic acids to assimilate NH_4_^+^ and produce energy for plant growth.

### Defense response

Nitrogen status is often related to plant disease emergence and plant immunity (Fagard et al., 2014). It is well known that NH_4_^+^ induces the upregulation of defense-related genes *in planta* (Ravazzolo et al., 2020). The results presented here demonstrate the transcriptional activation of defense-related genes, such as those encoding for class IV chitinase, defensin 5.1, beta-lactamase and a thaumatin-like protein, in maritime pine roots in response to the NH_4_^+^ supply (Supplemental Dataset 2). A similar tendency was observed at the proteomic level for several ginkbilobin antifungal proteins (Wang and Ng, 2000). This effect highlights the importance of the upregulation of this kind of genes in conifers in their natural environment since these plants have adapted to living in suboptimal environments where nutrient availability is a limiting factor for their growth and development (Farjon, 2018).

In this context, the upregulation of *antimicrobial peptide 1* and *pathogenesis-related protein PR-1* transcripts was consistent with previous results in maritime pine roots (Canales et al., 2010). Pine AMP1 is known to inhibit fungal development (Canales et al., 2011). It has been proposed that, related to its defensive function, AMP1 can inhibit the NH_4_ uptake in maritime pine, possibly through a physical interaction with ammonium transporters (AMT) (Castro-Rodríguez et al., 2017). Additionally, related to AMT activity regulation, transcriptomic results showed a wide repression of several transcripts coding for *calcineurin B-like* (*CBL)-interacting protein kinases* and *calcium-binding proteins* (Supplemental Dataset 2); in the second case, the transcript also contained significant modifications (Supplemental Dataset 10). The protein kinase CIPK23 has been shown to phosphorylate T460 of AtAMT1.1 and AtAMT1.2 in a CBL-1-dependent manner, repressing NH_4_^+^ transport (Straub et al., 2017). These results imply transcriptional regulation of NH_4_ ^+^ uptake resulting in proper N acquisition and assimilation. Accordingly, no statistically significant changes in the AMT proteins were observed, likely due to the relevance of their posttranslational regulation (Figure 3; Supplemental Dataset 2).

### Ethylene

Transcriptomic studies carried out in rice described that under excess NH_4_^+^, ethylene (ET) could be one of the major regulatory molecules in roots (Sun et al., 2017). Additionally, Li et al. (2013) described in *Arabidopsis* that shoot-supplied NH_4_^+^ promoted ET biosynthesis only in shoots, resulting in a reduction in the lateral root formation process due to *auxin transporter 1* (*AUX1*) repression. These data might indicate that ET biosynthesis is involved in detrimental NH_4_^+^-related phenotypes, such as a reduction in the number of lateral roots (LRs). Some reports have described that the application of inhibitors of the ET biosynthesis improved symptoms of NH_4_^+^ toxicity affecting RSA (Li et al., 2013). However, previous results indicated that NH_4_^+^ promotes root growth in maritime pine seedlings (Ortigosa et al., 2020).

A comparison of the root transcriptomic response and differential proteomics results revealed that NH_4_^+^ promotes a decrease in ET-related transcripts and proteins. Several ET-responsive TFs were downregulated at the transcriptomic level, while proteomic results revealed that ACC oxidase was downregulated (Supplemental Datasets 2 and 7). ACC oxidase is the enzyme responsible for the final stage in the biosynthesis of ET in plants (John et al., 1999). Furthermore, ACC oxidase transcripts exhibited a nucleoside putative modification in the 3’-UTR (position 462) that was differentially increased under NH_4_^+^ treatment (Supplemental Dataset 10). Since the levels of ACC oxidase transcripts had no significant changes, it is possible that this epitranscriptomic mark could be involved in the translational regulation of these transcripts, therefore affecting the ET levels of the organ. Interestingly, it is reasonable to think that this kind of response could be based on conifer preference for NH_4_^+^. Additional technical approaches are required to identify the exact modification and validate its biological relevance in the regulation of ACC oxidase and ET synthesis. Finally, the relationship between ET and N nutrition is remarked by the differential expression (negative) of a trihelix family TF that has a very high similarity degree to *AT3G54390* in *Arabidopsis* (Figure 7). This TF in *Arabidopsis* can interact with the promoters of genes involved in N metabolism and ET biosynthesis including *glutamine synthetase* (AT1G66200), *nitrite reductase* (AT2G15620), *ET-responsive transcription factor* (*ERF012*; AT1G21910) and *1-aminocyclopropane-1-carboxylate synthase* (AT5G65800) (Gaudinier et al., 2018).

**Figure 7.**
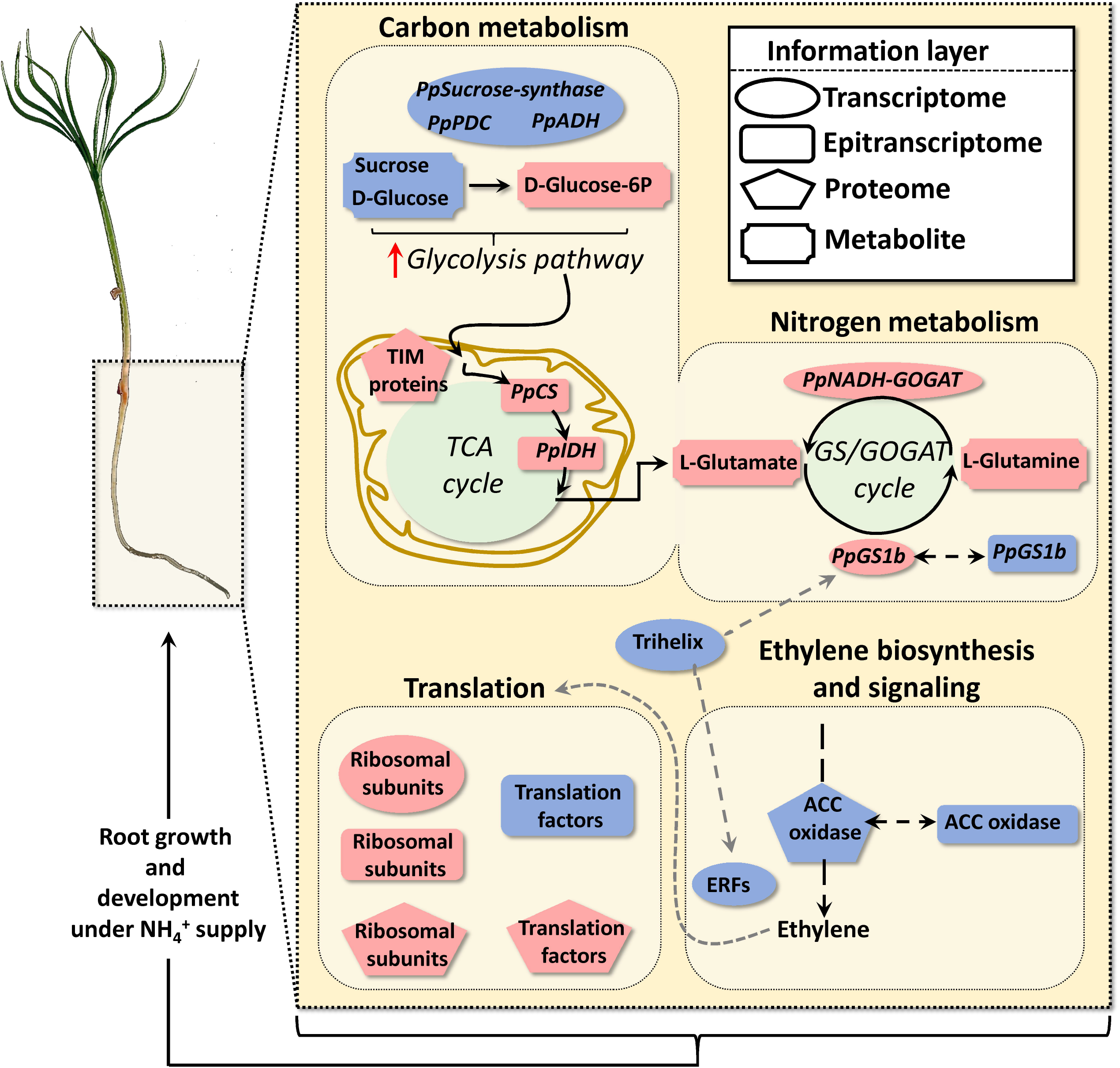
Schematic representation of functional results obtained through the omics integrative approach. NH_4_^+^ triggers carbon and nitrogen metabolism, ethylene biosynthesis and signaling and translation responses. Geometric red and blue forms indicate upregulation and downregulation respectively of transcripts, RNA nucleoside modifications, proteins, and metabolites.

### Translation and growth

The results also indicate a reconfiguration of the ribosomal proteins and elongation factors at all biological levels, although this was more visible in epitranscriptomics and proteomics results (Figure 6D and 7; Supplemental Datasets 2, 4, 7 and 10). This process is clearly related to the correlation between m^6^A ratios and protein amounts (Figure 5C), suggesting that NH_4_^+^ nutrition promotes general translation activation to support the root growth of maritime pine seedlings. This kind of effect on the ribosomal protein composition and proteins involved in translation has been previously observed under different conditions, including plant mineral nutrition (Wang et al., 2013; Prinsi and Espen, 2018; Ferretti and Karbstein, 2019). The data suggest that changes in the profiles of ribosomal proteins can be significantly mediated by epitranscriptomic modifications to their transcripts. These posttranscriptional changes can influence ribosome function and performance. One example is the RACK1 proteins, which are components of the small subunit of the ribosome and are involved in translation quality control promoting the endonucleolytic cleavage of nonstop mRNA (Ikeuchi and Inada, 2016). These proteins in plants are involved in the response to phytohormones, including ET, and in the regulation of growth and development (Wang et al., 2019). Interestingly, the mRNA of a *RACK1* gene in maritime pine roots has several differential epitranscriptomic marks (Supplemental Datasets 4 and 5). Although it was not significant, RACK1 protein expression was repressed under NH_4_^+^ conditions, suggesting a change in the quality control mechanisms and even in the integration of the plant signaling pathways. This result agrees with the negative regulation of ACC oxidase and repression of several ERFs (Supplemental Datasets 2 and 7). Accordingly, ET signaling affects gene translation in different ways (Merchante et al., 2015).

Additionally, related to the regulation of translation and root growth, among the transcriptomic results, the repression of an *FCS-like zinc finger* was observed (Supplemental Dataset 2). FCS proteins interact with SnRK1 to mediate the interaction of SnRK1 with other proteins (Jamsheer et al., 2018). SnRK1 is a kinase that phosphorylates RAPTOR, a regulatory element of the TOR complex, repressing TOR activity (Fu et al., 2020). Considering the ribosomal protein changes and the increase in root growth under NH_4_ nutrition in maritime pine seedlings (Ortigosa et al., 2020), it can be hypothesized a decrease of SnRK1 activity over the TOR complex in these conditions. The TOR complex integrates different developmental and environmental signals, including nutritional status, promoting ribosome biogenesis, translation, and plant growth.

Finally, a transcript coding for a TCTP protein had several differential modifications (Supplemental Dataset 5). It is well established in plants that *TCTP* mRNA is synthetized in shoots and transported to roots, where it is translated and promotes lateral root formation (Branco and Masle, 2019). Interestingly, transport of this mRNA is mediated through epitranscriptomic marks (Yang et al., 2019). This suggests additional mechanisms controlling growth and development in response to N nutrition in maritime pine roots.

Therefore, the results presented in this work suggest that protein translation and growth are finely regulated through epitranscriptomic marks, including m^6^A, to acquire an optimum response to N supply (Figure 7). More research efforts are required to corroborate this hypothesis and to investigate whether ET could act as a modulator of the integrated response observed due to its effects on the translation process.

## MATERIALS AND METHODS

### Plant material

Seeds from maritime pine (*Pinus pinaster* Ait.) from “Sierra Segura y Alcaraz” (Albacete, Spain) were provided by from the *Área de Recursos Genéticos Forestales* of the Spanish *Ministerio de Agricultura, Pesca y Alimentación*. Maritime pine seed germination was carried out following the protocol described in (Cañas et al., 2006). Seedlings were grown in vermiculite in plant growth chambers under 16 light photoperiod, a light intensity of 100☐ μmol ☐m^−2^ ☐s^−1^, constant temperature of 25 °C and watered twice a week with distilled water. One-month old maritime pine seedlings were used for this experiment. Pine seedlings were randomly subdivided into two different groups, relocated into forestall seedbeds and watered with 80 mL distilled water. After three days of acclimation, the control group was irrigated with 80 mL of water (C) and the experimental group with 80 mL of 3 mM NH_4_Cl. Root samples were collected at 24 hours post-irrigation and immediately frozen in liquid N. This experiment was carried out three independent times. The adequate development of each experiment was verified through the gene expression analysis by RT-qPCR of two control genes, *PpAMT1.3* and *PpAMP1* following previous results (Canales et al., 2011; Castro-Rodríguez et al., 2016).

### Metabolite profile

The metabolites for ^1^H-NMR analysis were extracted following the protocol previously described by Kruger et al. (2008). Two hundred milligrams of frozen powder were used for extractions. The ^1^H-NMR analyses were performed on an ASCEND 400 MHz NMR Spectrometer (Bruker, Billerica, MA, USA) in the Bionand Centre (Malaga, Spain). The 1D-^1^H-NMR spectrum for each sample was obtained as previously described by Cañas et al. (2015). Quantitative analysis of the NMR spectra was performed using LCModel software (Provencher, 1993) and a previously generated reference metabolite spectral library (Cañas et al., 2015). The internal reference was an electronically generated signal, ERETIC (Akoka et al., 1999). The metabolite amounts were determined at millimolar concentrations. The metabolite contents were analyzed with MetaboAnalyst 4.0 (Chong et al., 2018). Data were normalized using the quantile method, then log transformation and mean centered. MetaboAnalyst 4.0 was used to construct a Heatmap and perform a t-test with the metabolite data.

### Total RNA isolation

Total root RNA from maritime pine seedlings was isolated following the protocol described by Liao et al. (2004) and modified by Canales et al. (2012). The RNA concentration and purity were determined *via* spectrophotometry on a Nanodrop ND-1000 (Thermo Scientific, Waltham, MA, USA). Purity was determined through the 260/280 and 260/230 ratios. RNA quality was also determined in a Bioanalyzer 2100 (Agilent, Santa Clara, CA, USA). The concentration was verified with a Qubit 4 Fluorometer (Invitrogen, Paisley, UK) and Qubit RNA BR, Broad-Range, Assay Kit (Invitrogen, Paisley, UK).

### mRNA isolation and preparation

Samples with a RIN value > 7 were selected to mRNA isolation. The poly(A)-RNA isolation was performed using Dynabeads™ mRNA Purification Kit (Invitrogen, Paisley, UK) following the manufacturer’s instructions. This process was carried out twice per sample to avoid rRNA contamination. poly(A)-RNA quality was determined in a Bioanalyzer 2100 (Agilent, Santa Clara, CA, USA). The concentration was verified with a Qubit 4 Fluorometer (Invitrogen, Paisley, UK) and Qubit RNA HS, High Sensitivity, Assay Kit (Invitrogen, Paisley, UK).

### Direct RNA sequencing (DRS) and differential epitranscriptomic analysis

Nanopore libraries for DRS were prepared from 1.65 up to 2.18 µg of isolated poly(A)-RNA using the Nanopore Direct RNA Sequencing kit (SQK-RNA001, Oxford Nanopore Technologies, ONT, Oxford, UK) according to manufacturer’s instructions. The DRS libraries were loaded onto a R9.4 SpotON Flow Cells (Oxford Nanopore Technologies, Oxford, UK) and sequenced until complete nanopores depletion. Extended method descriptions are in the Supplemental Methods 1. Basecalling was carried out with ONT Guppy software (https://community.nanoporetech.com). The resultant reads were filtered by quality (Q>9). Read alignment was made with minimap2 software (Li, 2018) using root transcriptome of maritime pine as reference. The alignment parameters were adjusted for DRS (−*uf* and -*k14*). Differentially expressed (DE) transcripts were identified using the edgeR package for R, the transcripts were normalised by count per million mapped reads (cpm) and filtered (2 cpm in at least 2 samples) (Robinson et al., 2010). The transcripts with False Discovery Rate < 0.05 (FDR < 0.05) were considered as differentially expressed.

ONT-DRS reads were used for *de novo* modification detection with the TOMBO software (Stoiber et al., 2017). The total mapped reads per base and the number of modified bases in each position were obtained using the *text_output_browser_file* method with the options *coverage* and *fraction*. Only the positions with at least 50 mapped reads were employed for subsequent analyses. A Fischer exact test was carried out for each transcript position to determine the differential expression of modified bases among control and NH_4_^+^ supplied samples. The transcript positions with a *P-*value < 0.05 and an absolute logFC > 0.5 were considered as differentially modified.

*Transdecoder* software (https://github.com/TransDecoder/transdecoder.github.io) were used to determine modification positions and nucleobases in the transcripts of the reference transcriptome. Identification of m^6^A sites were carried out with the bioinformatic pipeline *Nanom6A* using default parameters (Gao et al., 2021). The length of poly(A) tails were determined using *Nanopolish* 0.11.1 software package (https://github.com/adbailey4/nanopolish) with the *polya* function.

### Functional annotation and enrichment analyses

The transcriptome was functional annotated with BLAST2GO (Götz et al., 2008) using DIAMOND software with *blastx* option (Buchfink et al., 2015) result against the NCBI’s plants-*nr* database (NCBI Resource Coordinators, 2016). Blast results were considered valid with e-value < 1.0E-6. Singular enrichment analysis (SEA) of the GO terms was made in the AGRIGO v2.0 web tool under standard parameters using as GO term reference the whole assembled transcriptome annotation (Tian et al., 2017). Representative enriched GO was determined using REVIGO (Supek et al., 2011).

### RT-qPCR

The cDNA synthesis was performed using 1 μg of total RNA and iScript™ cDNA Synthesis Kit (Bio-Rad, Hercules, CA, USA) following manufacturer’s instructions. The qPCR primers were designed following the MIQE guidelines (Bustin *et al*., 2009). The primers are listed in the Supplemental Table 2. qPCRs were carried out using 10 ng of cDNA and 0.4 mM of primers and 2X SsoFast^TM^ EvaGreen® Supermix (Bio-Rad, Hercules, CA, USA) in a total volume of 10 μL. Relative quantification of gene expression was performed using thermocycler CFX 384™ Real-Time System, Bio-Rad, Hercules, CA, USA). The qPCR program was: 3 min at 95 °C (1 cycle), 1 s at 95 °C and 5 s at 60 °C (50 cycles) and a melting curve from 60 to 95 °C, to generate the dissociation curve in order to confirm the specific amplification of each individual reaction. The analyses were carried out as described by Cañas et al. (2014) using the MAK3 model in the R package *qpcR* (Ritz and Spiess, 2008). Expression data were normalized to two reference genes, SKP1/ASK1 and SLAP that were previously tested for RT-qPCR experiments in maritime pine (Granados et al., 2016). The qPCR analyses were made with three biological replicates and three technical replicates per sample.

### Validation of differential deposition of m^6^A by RT-qPCR

The validation of the differential deposition of m^6^A in the transcripts were made using the SELECT method (Xiao et al., 2018). A differential cDNA synthesis was made per each transcript with 30 ng of total RNA. qPCR determinations were made with 2 μL of cDNA. The expression level of each transcript was determined in parallel by RT-qPCR and their result were used to normalize the SELECT results. The primers are listed in the Supplemental Table 2. Extended method description can be found in the Supplemental Methods 1.

### Differential proteomics analysis

The proteins were extracted following the protocol described by González Fernández et al. (2014). The extractions were carried out with 200 mg of sample. Protein content was determined using a commercially kit (Protein Assay Dye Reagent; Bio-Rad, CA, USA) and bovine serum albumin as a standard (Bradford, 1976). Protein extracts were cleaned-up in 1D SDS-PAGE at 10% polyacrilamyde as described in Valledor and Weckwerth, 2014. Protein bands were cut off, diced, and kept in water at 4°C until digestion.

Protein digestion and nLC-MS2 analysis were carried out in the Proteomics Facility at Research Support Central Service, University of Cordoba. Nano-LC was performed in a Dionex Ultimate 3000 nano UPLC (Thermo Scientific, Waltham, MA, USA) with a C18 75 μm x 50 Acclaim Pepmam column (Thermo Scientific, Waltham, MA, USA). Eluting peptide cations were converted to gas-phase ions by nano electrospray ionization and analyzed on a Thermo Orbitrap Fusion (Q-OT-qIT, Thermo Scientific) mass spectrometer operated in positive mode. Extended method descriptions are in the Supplemental Methods 1.

Root transcriptome from *Pinus pinaster* was translated into the six open reading frames with *transeq* tool (Madeira et al., 2019). The output peptides chains were filtered by length, deleting those less than 50 amino acids (Romero-Rodríguez et al., 2014). To reduce the redundancy of proteins in the database, CD-HIT-EST with a 99% identity filter was used (Fu et al., 2012). The raw data were processed using Proteome Discoverer (version 2.3, Thermo Scientific). MS2 spectra were searched with SEQUEST engine against the reference proteome database. *In silico* peptide lists were created using the followings settings: trypsin digestion, a maximum of two missed internal cleavage sites per peptide, precursor mass tolerance of 10 ppm and fragment mass tolerance of 0.75 Da per fragment ions. Only peptides with a high confidence (FDR ≤ 0.01) and minimum XCorr of 2 were selected. The identified proteins were filtered by a minimum of two different peptides. *Minora* algorithm was used to determine relative quantification. The protein identification with redundancy is considered by Proteome Discoverer and SEQUEST software. Proteins sharing peptides were grouped and all those groups without a unique peptide were removed. Only proteins detected in 5 of the 6 samples were considered for subsequent analyses. The resultant proteins were annotated using BLAST with *blastp* option against NCBI’s non-redundant database (Camacho et al., 2008) and BLAST2GO software (Götz et al., 2008). Proteins with the same sequence annotation and quantification profile across the samples was considered the same protein selecting the protein with the longest sequence. Normalized data from Proteome Discoverer software were used for a differential expression analysis using the edgeR package for R (Robinson et al., 2010). Only the proteins with *P*-value < 0.05 were considered as differentially expressed.

### Accession Numbers

The sequences of genes examined in this study can be found in Genbank under the following accession numbers: KC807909 (*PpAMT1.3*) and HM210085 (*PpAMP1*). The root transcriptome sequences consulted at https://fmp.uma.es/sequenceserver/ or can also be obtained on demand from the contact author. The DRS data have been deposited in the NCBI’s Gene Expression Omnibus (Edgar et al., 2002) and are accessible through GEO Series with the accession number GSE174830 (https://www.ncbi.nlm.nih.gov/geo/query/acc.cgi?acc=GSE174830). The mass spectrometry proteomics data have been deposited to the ProteomeXchange Consortium via the PRIDE partner repository (Vizcaino et al., 2013) with the dataset identifier PXD025331 and 10.6019/PXD025331.

## AUTHOR CONTRIBUTIONS

F.O. have performed the experiments; C.L.F. has performed the bioinformatic analyses; J.P.C. and F.R.C. have performed the statistical data analysis; F.O. and R.A.C. have wrote the manuscript; C.A. and F.M.C. made additional contributions and edited the manuscript. F.O. and R.A.C. have planned and designed the research. R.A.C, C.A.V., and F.M.C were the responsible of the funding acquisition.

## ACKNOWLEDGEMENTS

This work was funded by Spanish *Ministerio de Economía y Competitividad*, grants numbers BIO2015-73512-JIN MINECO/AEI/FEDER, UE, BIO2015-69285-R and RTI2018-094041-B-I00. FO was partially supported by a grant from the *Universidad de Málaga* (*Programa Operativo de Empleo Juvenil vía SNJG, UMAJI11, FEDER, FSE, Junta de Andalucía*).

## Supplemental Information

**Supplemental Table 1.** Direct RNA sequencing information for each sample run.

**Supplemental Table 2.** Primer list.

**Supplemental Methods 1.** Extended Materials and Methods.

**Supplemental Figure 1.** Heatmap of root metabolites. The analysis was done with Metaboanalyst using the Euclidean distance and Ward clustering algorithm. Class 0 (blue group) corresponds with control samples irrigated with distilled water. Class 3 (red group) corresponds with seedlings irrigated with 3 mM NH_4_^+^.

**Supplemental Figure 2.** Significant GO terms after SEA analysis using the DRS results.

**Supplemental Figure 3.** Scatter plot and correlations between the transcript amounts and the modification ratios detected using Tombo software for each nucleoside.

**Supplemental Figure 4.** Significant GO terms after SEA analysis using the epitranscriptomics results obtained from Tombo software.

**Supplemental Figure 5.** Scatter plot and correlations between the poly(A) length and the modification ratios detected using Tombo software for each nucleoside and global results.

**Supplemental Figure 6.** Significant GO terms after SEA analysis using the proteomics results.

**Supplemental Figure 7.** Scatter plot and correlations between the protein amounts and the modification ratios detected using Tombo software for each nucleoside and global results.

**Supplemental Figure 8.** Glutamine synthetase activity in maritime pine roots under ammonium nutrition. The results are the mean of three independent experiments using biological pools. Error bars correspond to SE. Enzymatic determinations were described in Supplemental Methods 1.

**Supplemental Dataset 1.** Normalized data from root metabolite profiling.

**Supplemental Dataset 2.** Whole root DRS differential expression results. DE transcripts upregulated are in highlighted red. DE transcripts downregulated are in highlighted blue.

**Supplemental Dataset 3.** SEA whole root DRS differential overexpressed transcripts results.

**Supplemental Dataset 4.** DRS differential modification results in pine roots under ammonium nutrition. DE transcripts overmodified are in highlighted red. DE transcripts undermodified are in highlighted blue.

**Supplemental Dataset 5.** Transcripts with upregulated and downregulated modifications at the same time.

**Supplemental Dataset 6.** SEA of transcripts with significant epitranscriptomic modifications in pine roots under ammonium nutrition.

**Supplemental Dataset 7.** Identified differential m^6^A positions coincident with DRS differential modification detected by Tombo software in the three close positions context. DE transcripts overmodified are in highlighted red. DE transcripts undermodified are in highlighted blue.

**Supplemental Dataset 8.** Whole root differential proteomics results. DE proteins upregulated are in highlighted red. DE proteins downregulated are in highlighted blue. **Supplemental Dataset 9.** SEA results of proteins overexpressed in pine roots under ammonium nutrition.

**Supplemental Dataset 10.** Elements with significant changes at least in two experimental approaches.

**Supplemental Dataset 11.** Commons significant GO terms between omics analyses.

