## Supplemental Table 1 for "Epitranscriptome changes triggered by ammonium nutrition regulate the proteome response of maritime pine roots"

**Table S1.** Direct RNA sequencing information for each sample run.

| **Sample** | **Length read (mean, bp)** | **Length read (median, bp)** | **Maximum length read (bp)** | **Maximum quality per sample** |
| --- | --- | --- | --- | --- |
| C1 | 1022.96 | 887 | 14299 | 27.94 |
| C2 | 1009.87 | 854 | 11465 | 29.37 |
| C3 | 1059.41 | 889 | 10808 | 30.20 |
| N1 | 1031.77 | 886 | 12834 | 14.32 |
| N2 | 907.56 | 768 | 10951 | 29.96 |
| N3 | 927.18 | 786 | 10298 | 30.50 |
| **Mean** | **993.13** | **845** | **11775.83** | **27.05** |
