## Supplemental Method 1 for "Epitranscriptome changes triggered by ammonium nutrition regulate the proteome response of maritime pine roots"

**Supplemental Method 1.** Extended Materials and Methods

**Direct RNA sequencing (DRS) and differential epitranscriptomic analysis**

Nanopore libraries for DRS were prepared from 1.65 up to 2.18 µg of isolated polyA-RNA using the nanopore Direct RNA Sequencing kit (SQK-RNA001, Oxford Nanopore Technologies, ONT, Oxford, UK) according to manufacturer’s instructions. The poly(T) adapter ligation to the polyA-RNA was performed using T4 DNA ligase (New England Biolabs, Ipswich, MA, USA) in the Quick Ligase reaction buffer (New England Biolabs) for 15 min at room temperature. First-strand cDNA synthesis was carried with SuperScript III Reverse Transcriptase (Thermo Fisher Scientific, Waltham, MA, USA) using the oligo(dT) adapter. The RNA-cDNA hybrid obtained was purified using Agencourt RNAClean XP magnetic beads (Beckman Coulter, Brea, CA, USA). The DRS adapter was ligated to polyA-RNA using T4 DNA ligase (New England Biolabs, Ipswich, MA, USA) in the Quick Ligase reaction buffer (New England Biolabs, Ipswich, MA, USA) for 15 min at room temperature followed by a second purification step using Agencourt beads (as described above). Finally, DRS libraries were loaded onto a R9.4 SpotON Flow Cells (Oxford Nanopore Technologies, Oxford, UK) and sequenced until complete nanopores depletion, around 24 hours of run time.

**Validation of differential deposition of m^6^A by RT-qPCR**

The validation of the differential deposition of m^6^A in the transcripts were made using the SELECT method (Xiao et al., 2018). A differential cDNA synthesis was made per each transcript with 30 ng of total RNA in a reaction mix with 40 nM up-primer, 40 nM down-primer, 5 μM dTTP and 1x CutSmart buffer (New England Biolabs, Ipswich, MA, USA) in a final volume of 17 μL. The primers hybridization was made in a thermocycler at 90ºC for 1 min, 80ºC for 1 min, 70ºC for 1 min, 60ºC for 1 min, 60ºC for 1 min, 50ºC for 1 min and 40ºC for 6 min. After, 3 μL of a new solution containing 0.01 U BST2.0 DNA polymerase (New England Biolabs, Ipswich, MA, USA), 0.5 U SplintR ligase (New England Biolabs, Ipswich, MA, USA) and 10 nmol ATP was added to the reaction mix. The final solution was incubated in a thermocycler at 40ºC for 20 min with a final denaturation step at 80ºC for 20 min. Finally, 2 μL of the mix were employed to make qPCR determinations as described above. The expression level of each transcript was determined in parallel by RT-qPCR and their result were used to normalize the SELECT results.

**Differential proteomics analysis**

*Protein extraction*

The proteins were extracted following the protocol described by González Fernández et al. (2014). The extractions were carried out with 200 mg of sample. Protein content was determined using a commercially kit (Protein Assay Dye Reagent; Bio-Rad, CA, USA) and bovine serum albumin as a standard (Bradford, 1976). Protein extracts were cleaned-up in 1D SDS-PAGE at 10% polyacrilamyde as described in Valledor and Weckwerth, 2014. Protein bands were cut off, diced, and kept in water at 4ºC until digestion.

*Protein digestion*

Protein digestion were carried out in the Proteomics Facility at Research Support Central Service, University of Cordoba. Briefly, gel dices were distained in 200 mM ammonium bicarbonate (AB)/50% acetonitrile for 15 min followed by 5 min in 100 % acetonitrile. Protein was reduced by addition of 20 mM dithiothreitol in 25 mM AB and incubated for 20 min at 55 °C. The mixture was cooled down to room temperature, followed by alkylation of free thiols by addition of 40 mM iodoacetamide in 25 mM AB, in the dark for 20 min and the gel pieces were then washed twice in 25 mM AB. Proteolytic digestion was performed by addition of Trypsin (Promega, Madison, WI) at 12.5 ng/µL of enzyme in 25 mM AB and incubated at 37 ºC overnight. Protein digestion was stopped by addition of trifluoroacetic acid at 1% final concentration and the digested samples were finally Speedvac dried.

*nLC-MS2 analysis*

Protein analysis were carried out in the Proteomics Facility at Research Support Central Service, University of Cordoba. Nano-LC was performed in a Dionex Ultimate 3000 nano UPLC (Thermo Scientific) with a C18 75 μm x 50 Acclaim Pepmam column (Thermo Scientific). The peptide mix was previously loaded on a 300 μm x 5 mm Acclaim Pepmap precolumn (Thermo Scientific) in 2% acetonitrile/0.05% TFA for 5 min at 5µL/min. Peptide separation was performed at 40°C for all runs. Mobile phase buffer A was composed of water, 0.1% formic acid. Mobile phase B was composed of 20% acetonitrile, 0.1% formic acid. Samples were separated at 300 nL/min. Elution conditions were: 4-35%B for 60 min; 35-55% B for 3 min; 55-90% B for 3 min followed by 8 min wash at 90% B and a 12 min re-equilibration at 4%B. Total time of chromatography was 85 min.

Eluting peptide cations were converted to gas-phase ions by nano electrospray ionization and analysed on a Thermo Orbitrap Fusion (Q-OT-qIT, Thermo Scientific) mass spectrometer operated in positive mode. Survey scans of peptide precursors from 400 to 1500 m/z were performed at 120K resolution (at 200 m/z) with a 4 × 105 ion count target. Tandem MS was performed by isolation at 1.2 Da with the quadrupole, CID fragmentation with normalized collision energy of 35, and rapid scan MS analysis in the ion trap. The AGC ion count target was set to 2 x 103 and the max injection time was 300 ms. Only those precursors with charge state 2–5 were sampled for MS2. The dynamic exclusion duration was set to 15 s with a 10 ppm tolerance around the selected precursor and its isotopes. Monoisotopic precursor selection was turned on. The instrument was run in top 30 mode with 3 s cycles, meaning the instrument would continuously perform MS2 events until a maximum of top 30 non-excluded precursors or 3s, whichever is shorter.

**Glutamine synthetase enzyme activity**

Soluble proteins were extracted using 100 mg of root ground powder. The extraction was performed by adding 0.5 mL of extraction buffer (50 mM Tris-HCl pH 8, 1 mM EDTA, 10 mM MgCl2, 0.5 mM dithiothreitol (DTT), 20% (w/v) glycerol, 0.1% (v/v) Triton X-100, 1% (w/v) polyvinylpyrrolidone (PVP), and 1% (w/v) polyvinyl(poly)pyrrolidone (PVPP)) and 30 mg of fine sea sand. The resulting extract was centrifuged at 12,000 g for 30 min at 4 °C. The obtained supernatants were recovered and used for soluble protein determination through Bradford’s procedure using a commercial reagent (Protein Assay Dye Reagent; Bio-Rad, CA, USA) and bovine serum albumin as a standard (Bradford, 1976).

Glutamine synthetase (GS, EC 6.3.1.2) activity was determined by the transferase assay (Cánovas et al., 1984). The final reaction volume was 150 µL. Reactions were incubated for 15 min at 37 °C with 10 s of agitation every minute; the reactions were stopped with 150 µL of STOP solution (10% FeCl3·6H2O in 0.2 N HCl, 24% trichloroacetic acid and 50% HCl) and centrifuged for 3 min at 3220 g. After centrifugation, 200 µL of the supernatant was recovered, and absorbance was measured at 540 nm in a microplate reader.
