## Supplementary figures and images for "Epitranscriptome changes triggered by ammonium nutrition regulate the proteome response of maritime pine roots"

### Supplemental Figure 1

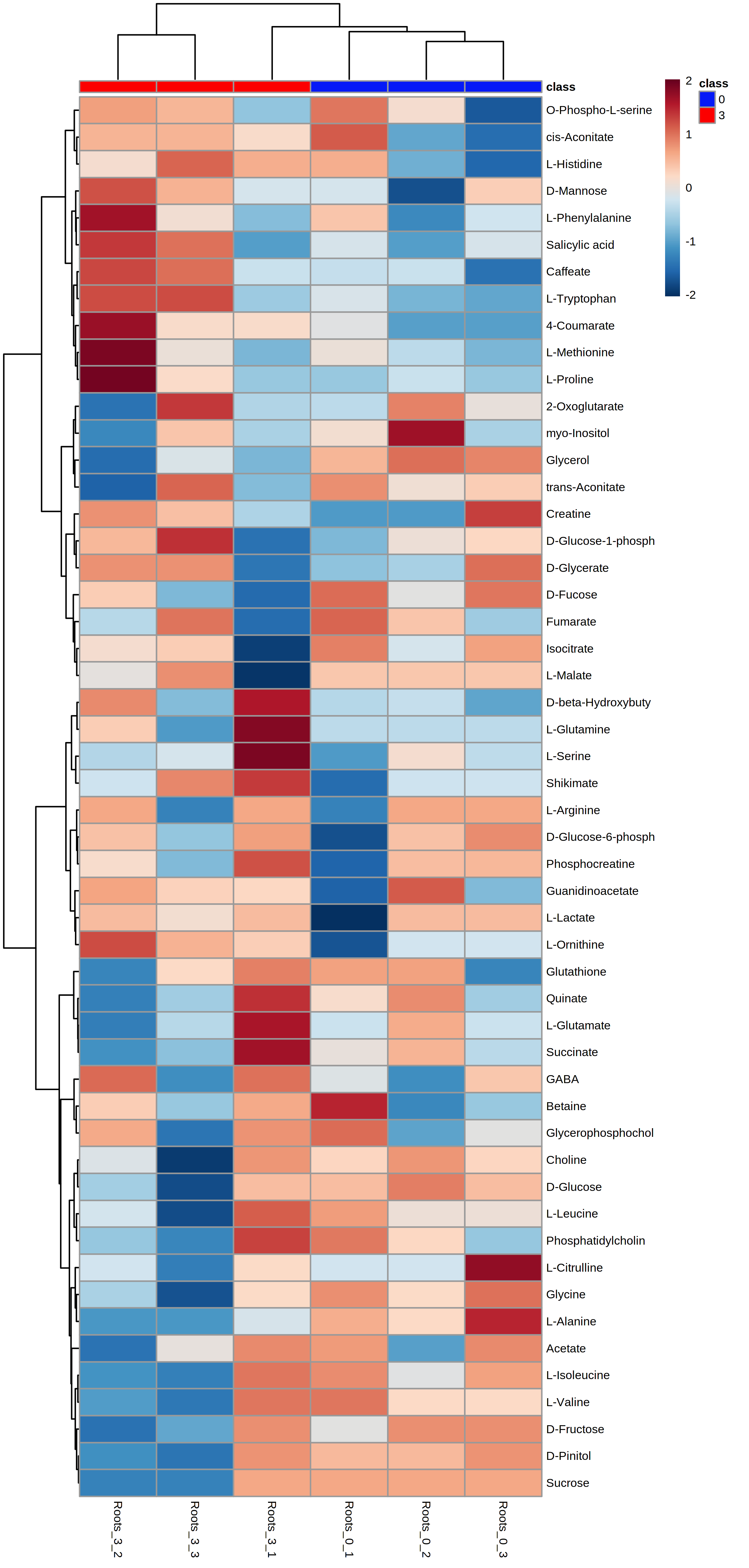

### Supplemental Figure 2

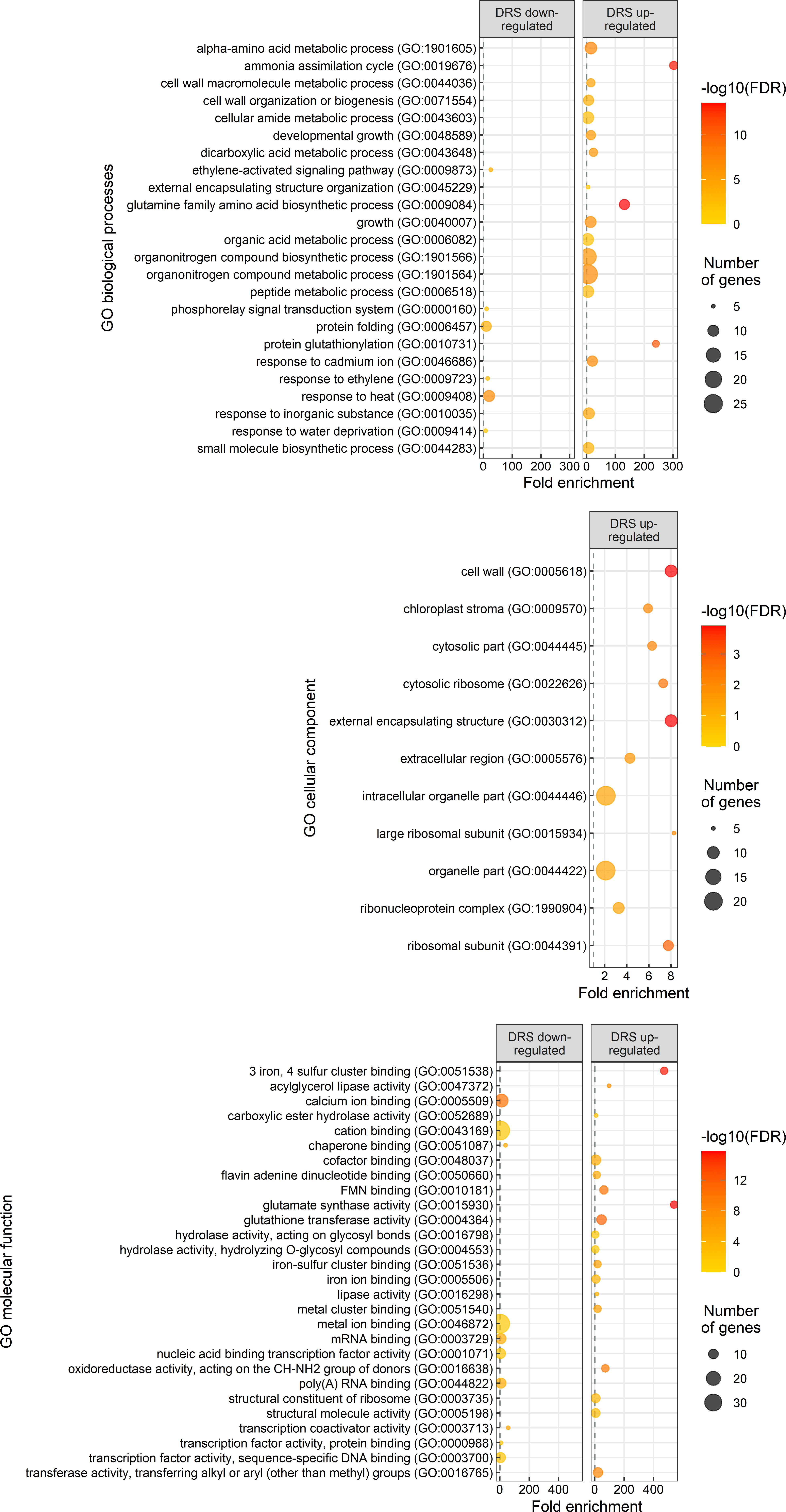

### Supplemental Figure 3

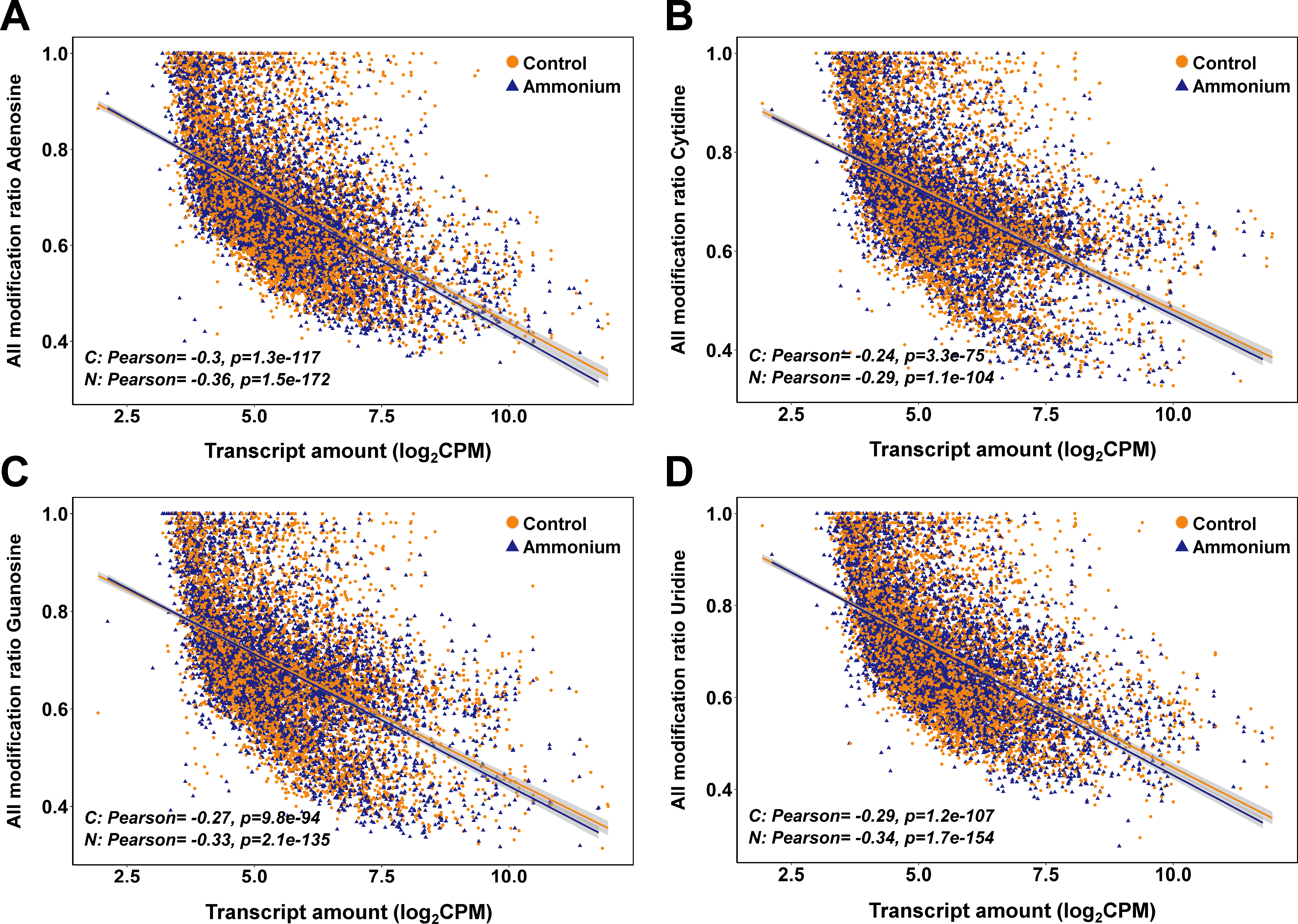

### Supplemental Figure 4

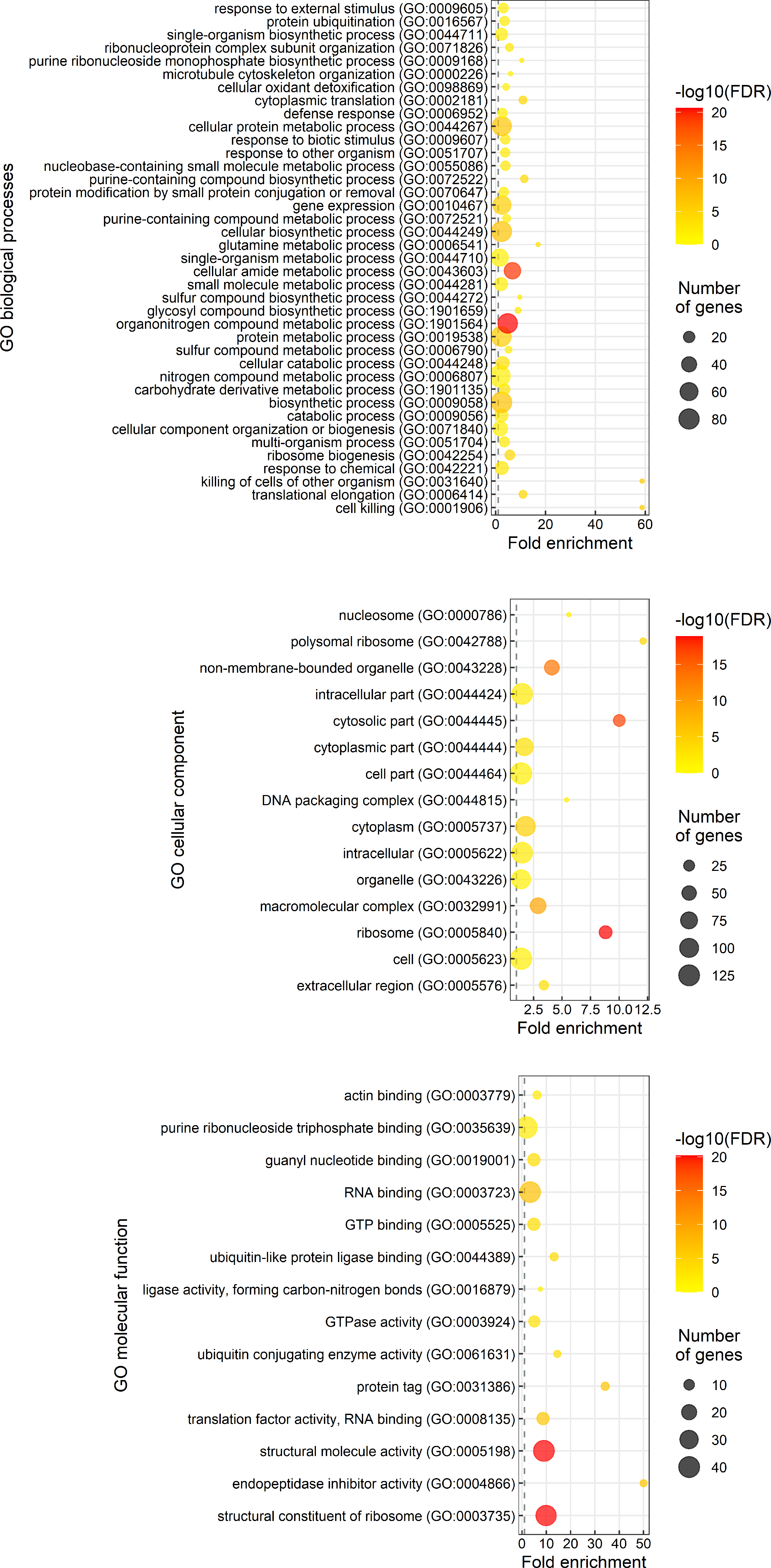

### Supplemental Figure 5

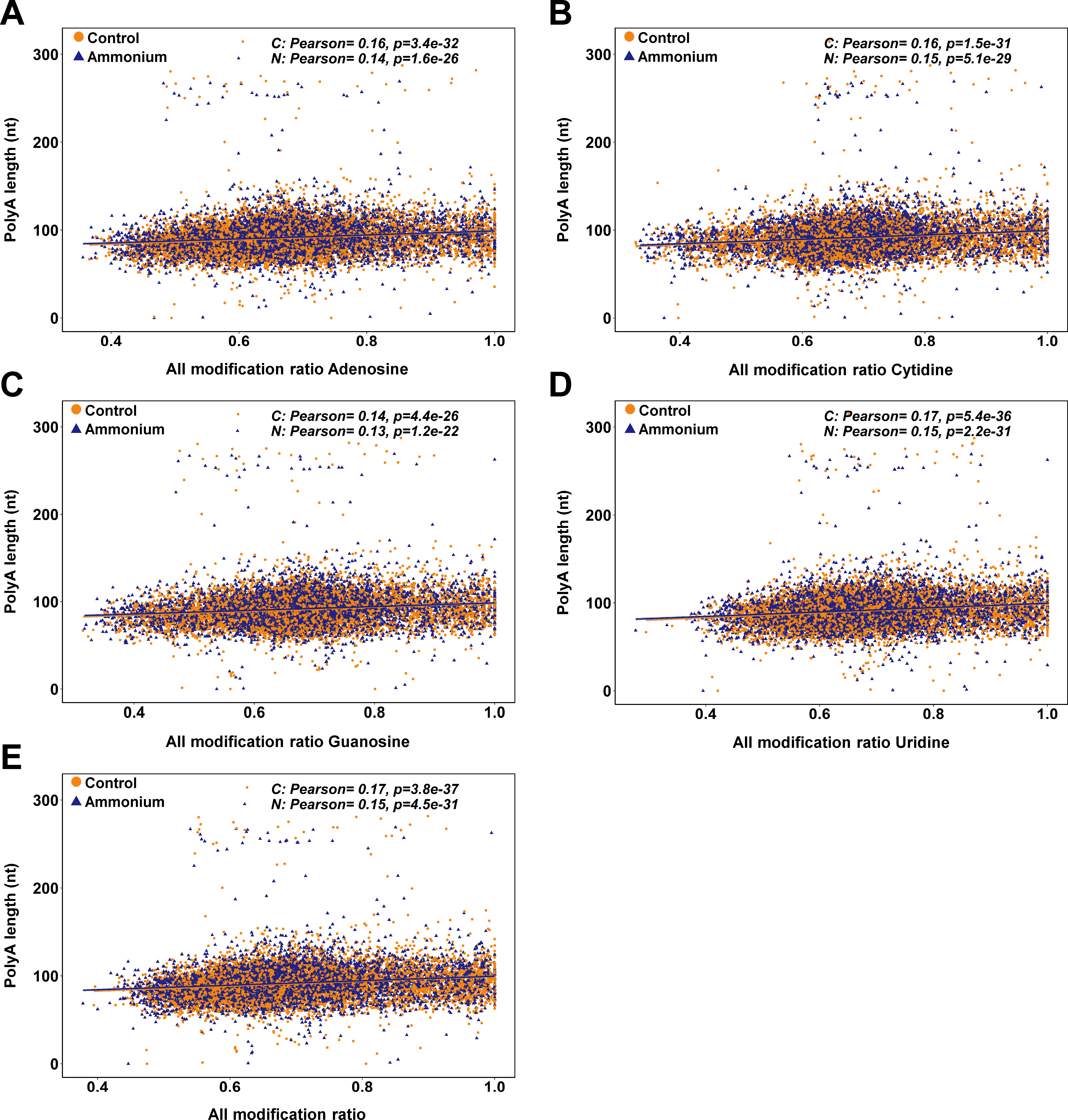

### Supplemental Figure 6

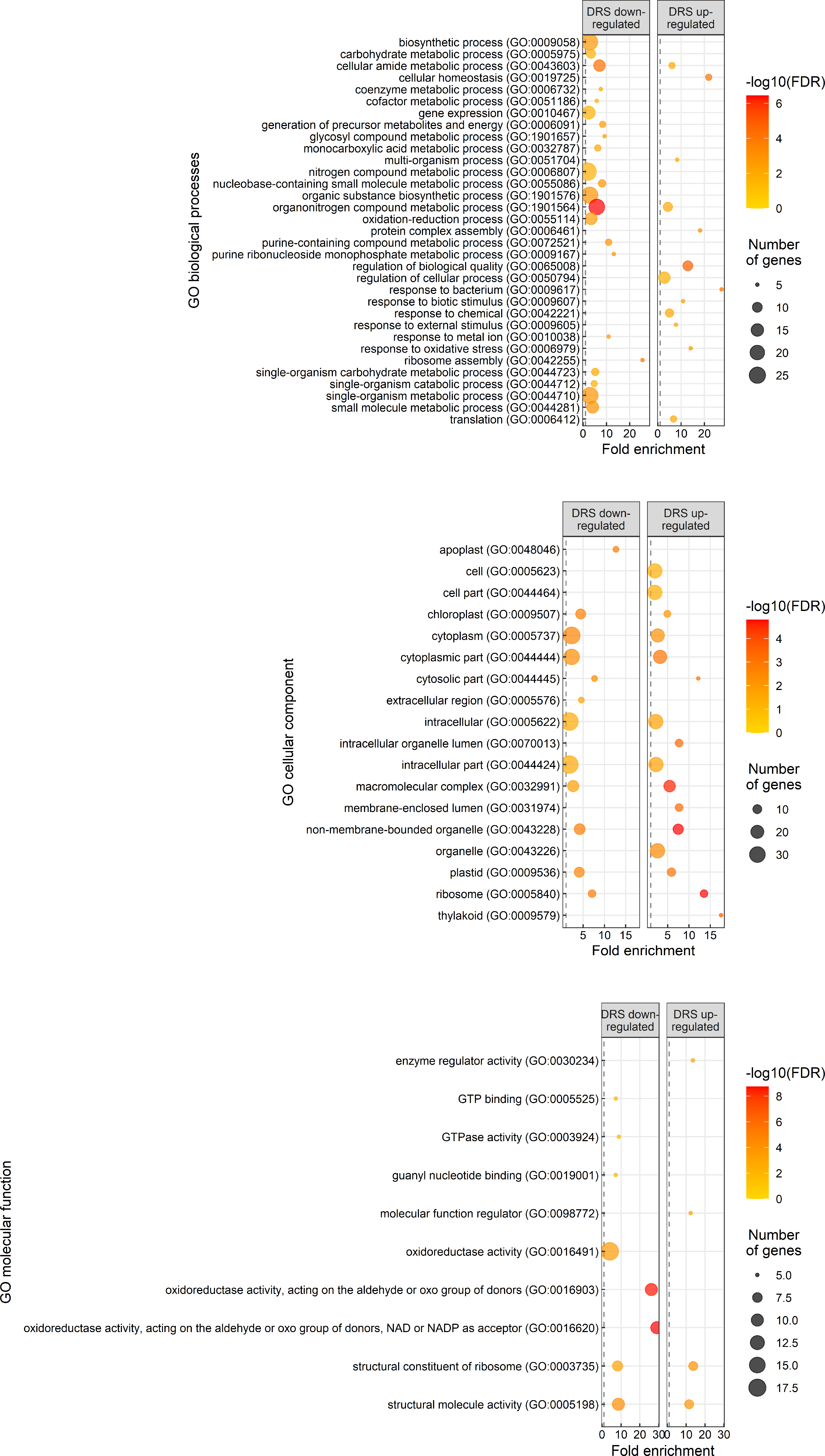

### Supplemental Figure 7

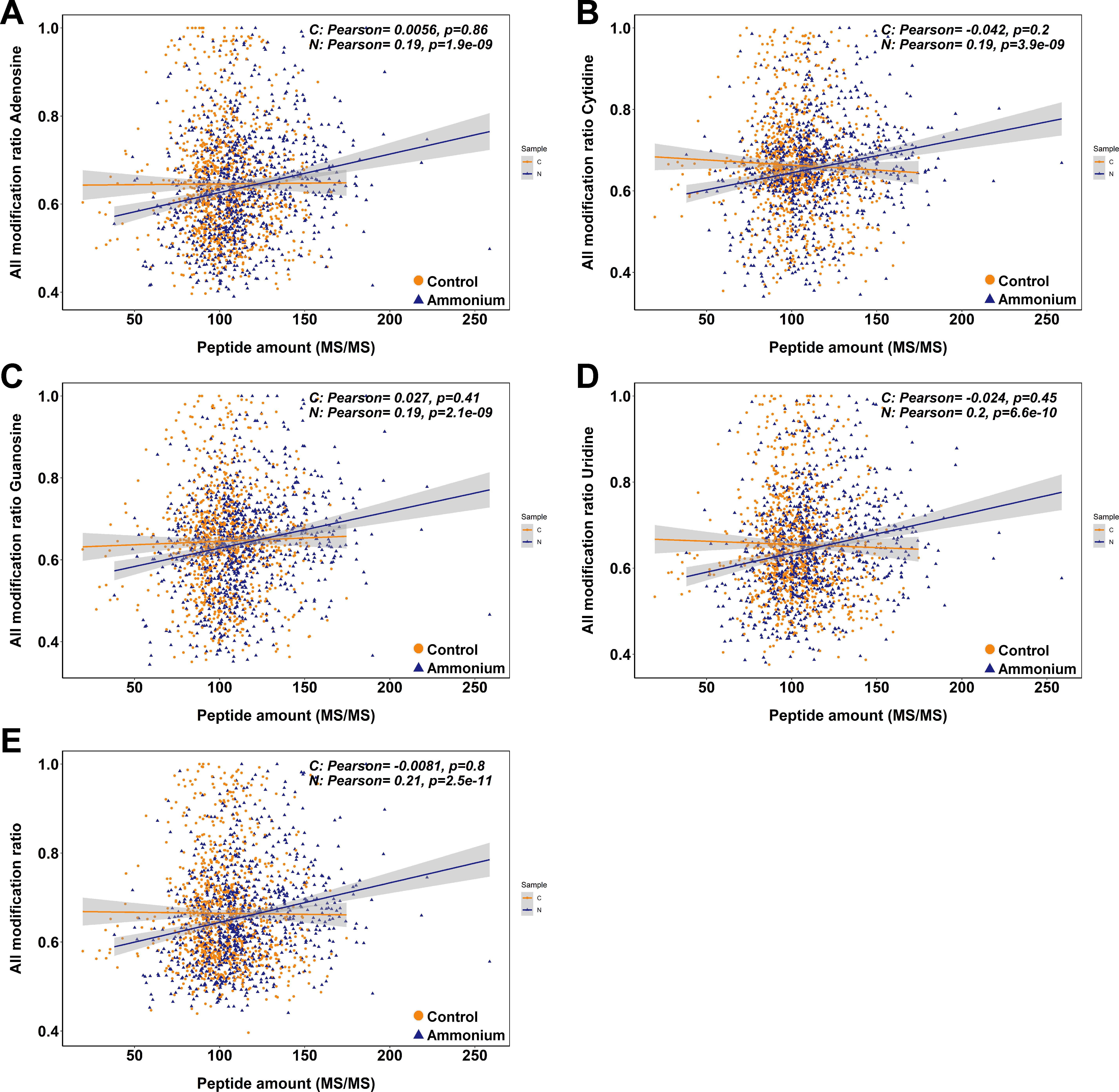

### Supplemental Figure 8

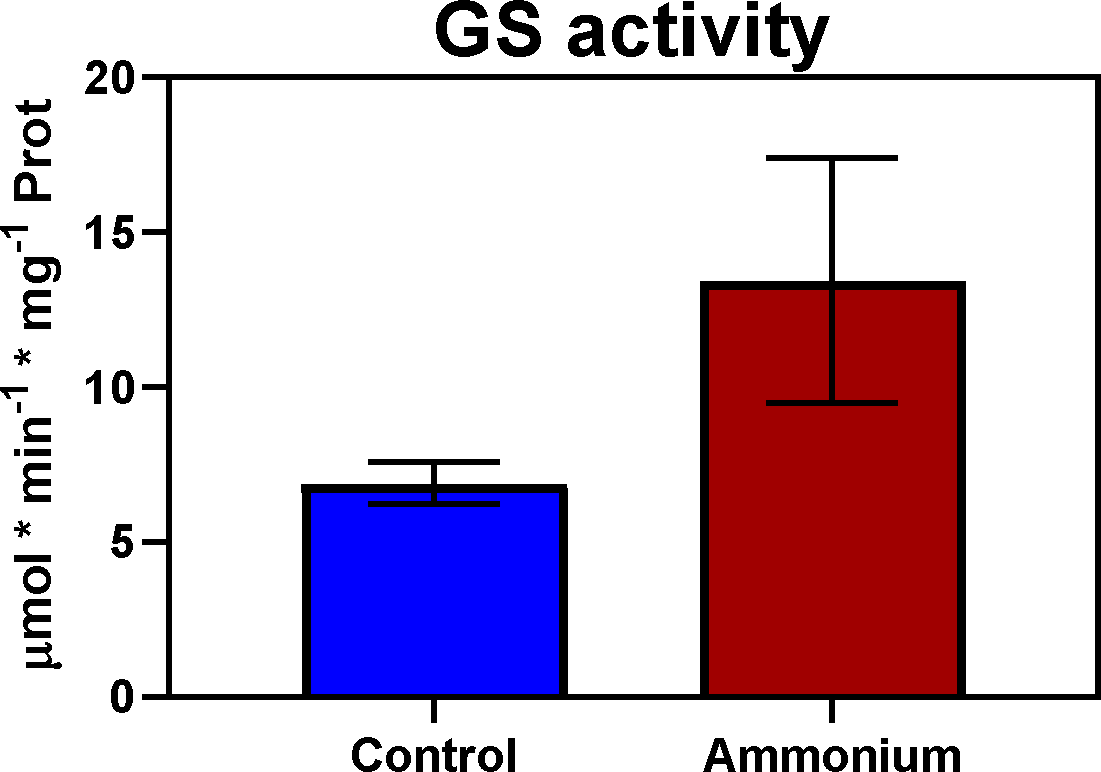
